# Metabolism-oriented compound screen in physiological culture conditions identifies a NAMPT inhibitor highly effective against drug-naïve and -resistant melanoma cells

**DOI:** 10.1101/2024.10.23.619802

**Authors:** Jasmin Renate Preis, Catherine Rolvering, Mélanie Kirchmeyer, Stephanie Kreis, Iris Behrmann, Claude Haan

**Affiliations:** Department of Life Sciences and Medicine, University of Luxembourg, Belvaux, Luxembourg

**Keywords:** melanoma, physiologic medium, melanoma 3D models, drug resistance, drug screening, FK866

## Abstract

Many melanoma patients do not respond to therapy or rapidly develop resistance to MAPK pathway inhibitors and immune checkpoint blockade treatments, highlighting the urgent need for additional therapeutic strategies for these patients.

To identify compounds that target drug-naïve and -resistant melanoma cells (Encorafenib/Binimetinib-resistant), we performed a screen using a metabolism-oriented compound library in different cell culture media and growth conditions (2D and 3D).

The efficacy of several compounds varied considerably under changing test conditions, but importantly, we also detected compounds that work across all tested conditions. In general, drug activity was reduced under hypoxia and in 3D spheroids. Thorough validation was performed for drugs showing potency in all conditions with a focus on efficacy in 3D spheroids grown in an in-house physiological culture medium. Using hydrogel matrix-embedded multi-cell type 3D models, we found that FK866, a nicotinamide phosphoribosyltransferase (NAMPT) inhibitor, is very potently suppressing melanoma cell growth while not affecting the growth of healthy cells such as fibroblasts and endothelial cells.

## Introduction

Melanoma is the most aggressive form of skin cancer with an increasing incidence over the last decades. Especially, once metastasized, melanoma is associated with poor patient survival (Whiteman et al., 2016). Mutations in genes encoding components of the mitogen-activated protein kinase (MAPK) pathway are found in most melanomas, with *BRAF* mutations being present in approximately 50% of patients (Ascierto et al., 2024; Colombino et al., 2012; Czarnecka et al., 2020; Lelliott et al., 2021; Winder & Virós, 2018). Consequently, BRAF and MEK inhibition has become the standard of care for patients with BRAFV600E mutations. Notably, the 5-year overall survival rate of BRAFV600F positive melanoma has increased from less than 10% to 35% for patients receiving combined Encorafenib (BRAFi) and Binimetinib (MEKi) treatment (Dummer, 2022). Furthermore, the 4-year overall survival rate has increased to 65% with the combination of immunotherapy and Encorafenib/Binimetinib (Ascierto et al., 2024). However, still many patients do not respond to therapy or rapidly develop resistance to MAPK pathway inhibitors and immune checkpoint blockade treatments (Winder & Virós, 2018), highlighting the urgent need for new therapeutic strategies. Many different mechanisms conferring kinase inhibitor resistance have been described (Kozar et al., 2019), which include metabolic rewiring and high metabolic plasticity of melanoma cells (Alkaraki et al., 2021; Avagliano et al., 2020).

Targeted compounds identified in classic preclinical *in vitro* 2D cell cultures regularly perform poorly in *in vivo* models or in clinical trials. An increasing number of studies shows that nutrient availability and concentrations can influence cancer cell sensitivity towards drugs (Cantor et al., 2017; Muir et al., 2017, 2018) in cell culture models. Along the same lines, using 3D cell culture models for drug testing have demonstrated a more faithful prediction of tumour drug responses (Lee et al., 2018; Vlachogiannis et al., 2018) and the metabolic phenotype of spheroid cultures grown in a physiological medium was described to better approximate the metabolome of *in vivo* tumours (Vande Voorde et al., 2019).

To apply these new findings to tackle melanoma resistance, we performed a drug screening with a metabolically targeted library in more physiological conditions: 3D cell culture models using drug sensitive and drug-resistant (DR) melanoma cells grown in an in-house cell culture medium (MPM). For this purpose, we established melanoma cell lines resistant to Encorafenib and Binimetinib by prolonged exposure to both drugs. Furthermore, we established a more physiological culture medium (MPM: modular physiologic medium) (Preis et al., 2024) better recapitulating the plasma concentration of metabolites. The screen was designed to show differences in compound activity for drug-sensitive and drug-resistant cells under the following conditions: (i) RPMI *vs*. MPM medium in 2D cultures (normoxia only) to identify drugs which act differently in the two media (different nutrient levels and FBS concentration), (ii) normoxia *vs*. hypoxia in 2D cultures (MPM only) to identify drugs changing efficacy in hypoxia, (iii) 2D cultures (normoxia and hypoxia) *vs*. 3D using the MPM medium to identify drugs which are influenced by nutrient gradients and/or perform differently due to an altered penetrance into 3D spheroids. With this multi-parametric approach (Figure 1) we identified compounds whose efficacy is environment-dependent, but also compounds that work across all tested conditions. Most of the metabolism targeting drugs had to be used in concentrations above 10 micromolar, indicating that for many metabolic targets the development of more potent compounds would be of advantage. Importantly, two inhibitors previously not connected to melanoma growth inhibition were identified to potently suppress cell viability of drug-resistant melanoma at low nanomolar concentrations: Torin2, an mTOR inhibitor and FK866, a NAMPT inhibitor.

**Figure 1.**
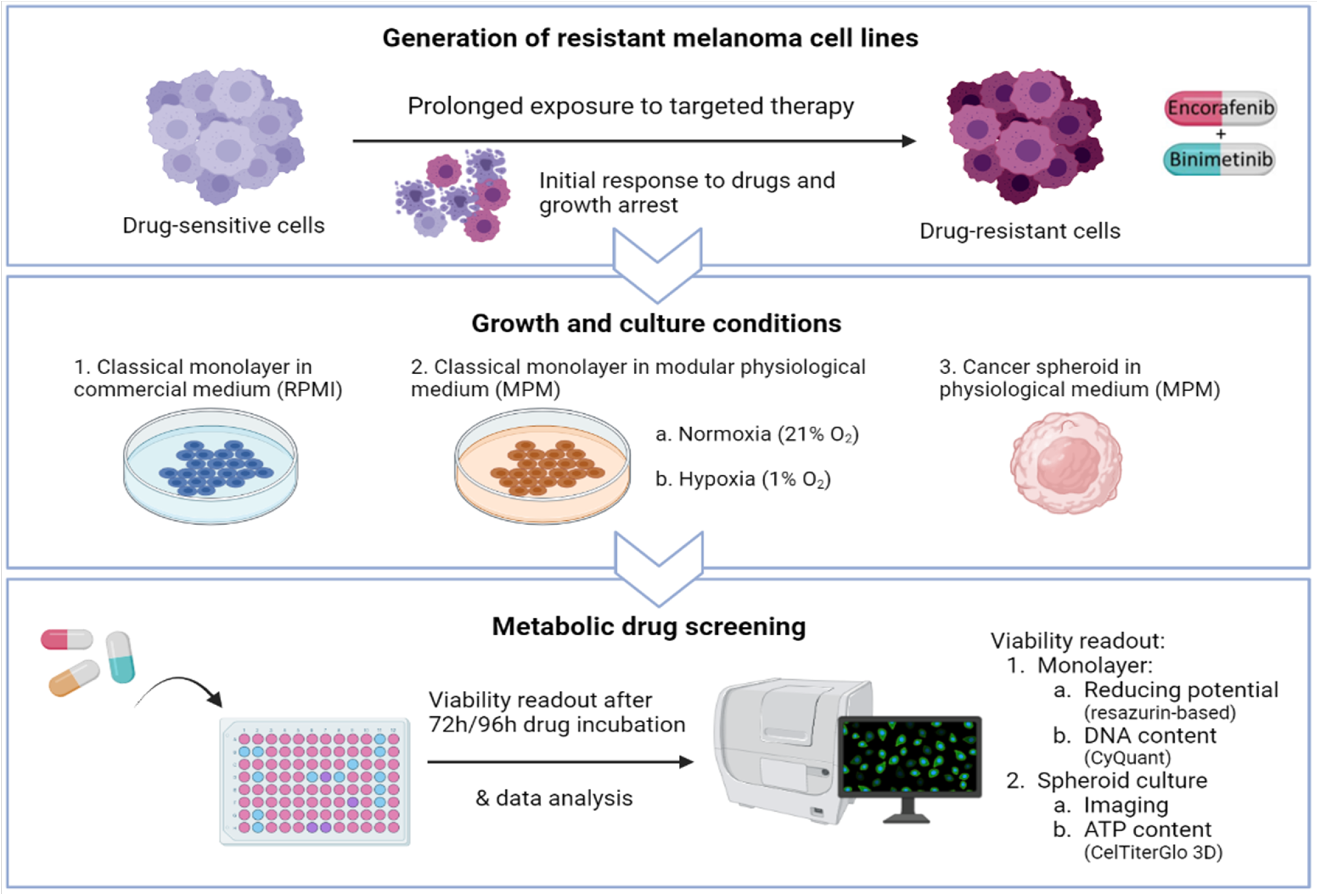
**Scheme depicting the screening approach**: Top panel: generation of drug-resistant melanoma cell lines by incubation with Encorafenib and Binimetinib for aprolonged period. Middle panel: culture conditions used to perform the screen and parts of the validations. Lower panel: overview of the screen and the readout assays used for screening.

## Materials and methods

### Cell culture and materials

All cells were grown at 37°C in a water-saturated atmosphere at 5% CO_2_. All BRAF-mutated melanoma cell lines (624Mel (Dr. Ruth Halaban), A375 (ATCC), WM3248 (Rockland)) were maintained in RPMI (Gibco) supplemented with 10% FBS. For experiments in modular physiologic medium (MPM) the cells were passaged and adapted for at least two days and then seeded for experiments. In short, MPM (see Preis et al., 2024, for the exact composition) is based on EBSS, contains 2.5% dialysed FBS and the BME vitamin mix. Physiological glucose (5 mM), amino acids (3.1 mM in total) and water-soluble metabolites (6,7 mM in total) found in human plasma constitute 34%, 21% and 45% of the nutrient concentration in MPM. MPM is similar to the physiologic media HPLM and Plasmax that have been described before (Cantor et al., 2017; Vande Voorde et al., 2019). For 3D spheroid cultures and in Organoplates (Mimetas) , MPM was supplemented with xeno-free B27 supplement (Gibco) and with growth factors (Insulin 500 ng/mL, HGF 50 ng/mL, BMP4 20 ng/mL, EGF 5 ng/mL, Hydrocortisone 1 µg/mL) to better mimic the physiologic situation (this modified version is called Mel-MPM) (see Preis et al., 2024 for the exact composition).

HMEC-1 (ATCC) were maintained in MCDB131 (Gibco) supplemented with 10 ng/mL Epidermal Growth Factor (EGF, Peprotech), 1 µg/mL Hydrocortisone (Merck), 10 mM Glutamine (Gibco) and 10% FBS (Gibco). NHDF (PromoCell) were cultivated in DMEM (Gibco) supplemented with 25 mM HEPES (Gibco) and 10% FBS (Gibco). Adaptation of NHDF and HMEC-1 to Mel-MPM was performed 7-10 days before starting the experiments. HMEC-1 cells were transduced with pLV-Bsd-CMV-tagBFP (Bio-connect) and selected with 10 µg/mL of blasticidin to yield the HMEC1-BFP. NHDF-iRFP were generated by transduction with LV-iRFP713-P2A-Puro (Imanis) and selected with 1 µg/mL puromycin. Both cell lines were sorted on a FACS Aria (BD biosciences).

### Generation of drug-resistant (DR) melanoma cell lines

The selected melanoma cell lines (624Mel, A375, WM3248) were subjected to prolonged exposure with the BRAF inhibitor Encorafenib and the MEK inhibitor Binimetinib (MedKoo Biosciences). The concentrations used to generate drug-resistant (DR) cells correspond to approximatively 10 times the respective IC_50_ concentrations of both individual drugs (Table in Figure 2B). The exposure was continued until the cells re-proliferated in the presence of the drugs, which took up to several months, depending on the cell line. The resistant cell lines were continuously exposed to Encorafenib and Binimetinib during passaging. Drug-naïve and drug-resistant cell lines were authenticated by STR profiling. WM3248-DR and 624Mel-DR cells were transduced with firefly luciferase-GFP lentivirus (CMV, Puro; Cellomics) and selected with 0.5 or 0.75 µg/mL puromycin to yield the WM3248-DR-GFP/luc and 624Mel-DR-GFP/luc cells, respectively. As we do not use luciferase assays in this publication, the cells are referred to as WM3248-DR-GFP and 624Mel-DR-GFP in the following text. GFP-positive cells were sorted on a FACS Aria (BD biosciences).

**Figure 2.**
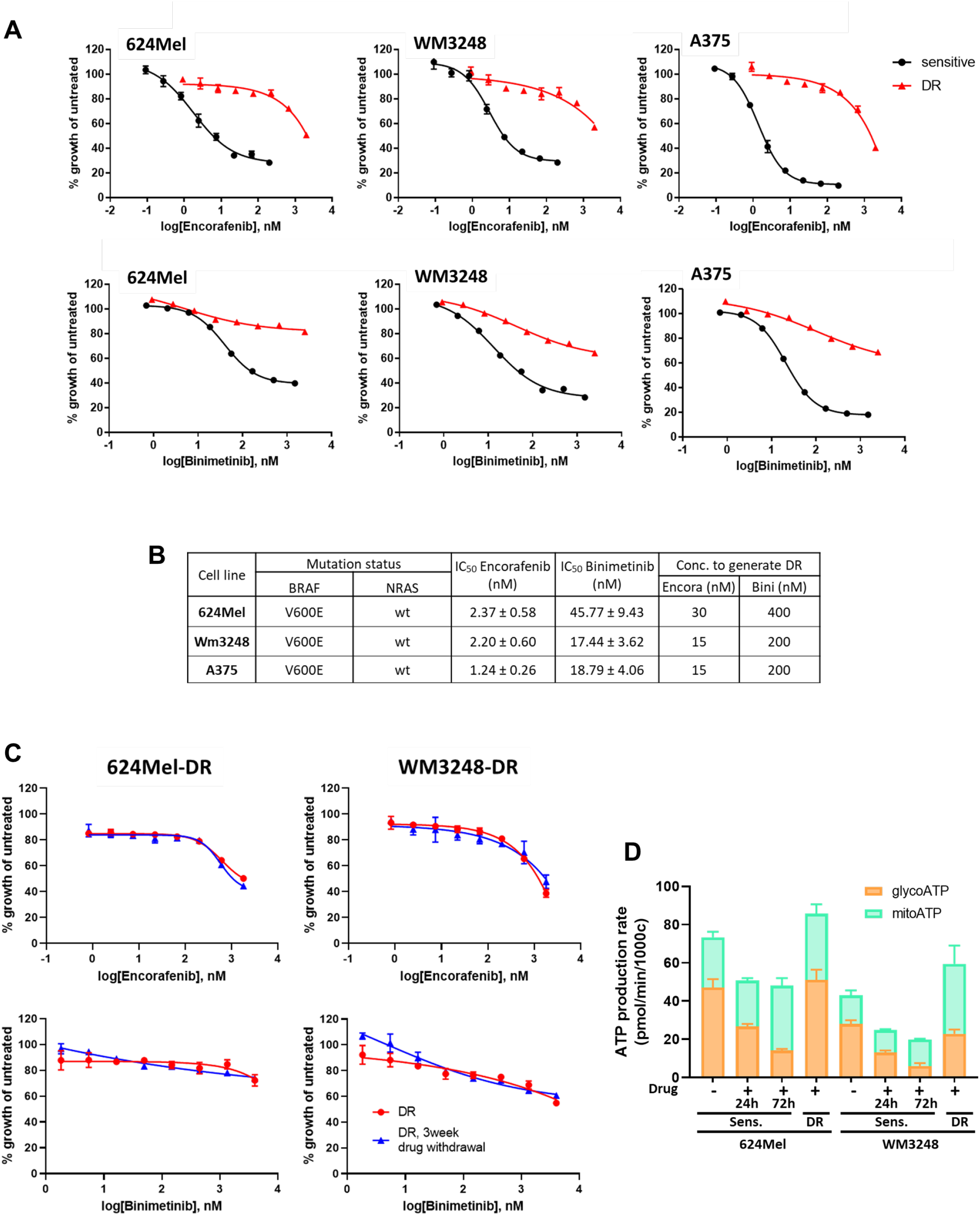
Generation and characterisation of drug-resistant melanoma cells. A) Dose-response curves for Encorafenib and Binimetinib for three different drug-sensitive cell lines (black curves) and the drug-resistant (DR) cell lines generated from them (red curves). 8 dilutions (1:3) per drug were used at starting concentrations of 200 nM Encorafenib and 1.5 µM Binimetinb for sensitive cells, and 2 µM Encorafenib and 2.5 µM Binimetinib for drug-resistant (DR) cells. Mean and standard deviation of one representative biological replicate out of 3 are shown, each performed in 3 technical replicates. B) Information on drug sensitive melanoma cell lines, their IC_50_ values to Encorafenib and Binimetinib and concentrations used to generate DR cells. C) Drug-resistant melanoma cells from which Encorafenib/Binimetinib was withdrawn for 3 weeks or 3 days were treated as described in A). Mean and standard deviation of one representative biological replicate out of 3 are shown, each performed in 3 technical replicates. D) ATP production rates of sensitive 624Mel and WM3248 cells with or without Encorafenib and Binimetinib treatment (24 h or 72 h) and in DR cells. The overall ATP production rate comprises ATP produced from glycolysis (glycoATP, orange) and from mitochondrial respiration (mitoATP, green), n=3 biological replicates, error bars represent mean ± SD.

### 2D and 3D compound screen

42 compounds targeting a variety of metabolic pathways and the individual drug concentrations (C1 and C2, C2 being 10 times higher than C1) were selected based on literature research (Figure S1 and Table S1). For the screen, the glucose concentration in RPMI was lowered to that of MPM (5 mM) to prevent that the differences seen for the metabolism-targeting compounds were due to the different glucose levels. The 2D screen was performed in technical duplicates. Each plate included 42 compounds and 6 positive and negative controls for the different readouts (medium only, untreated cells, cells treated with DMSO; cells treated with 500 nM Encorafenib, with 200 µM Etoposide, or with 1% Triton). Cells were seeded in black-walled 96-well µclear plates (Greiner Bio-one) for 2D adherent cell growth. After 3 days of drug treatment, cell growth was measured using the CyQuant Direct Cell Proliferation Assay Kit and PrestoBlue (both Life Technologies) with a parallel detection protocol. Fluorescence of CyQuant and PrestoBlue were measured on the Cytation 5 plate reader (Agilent Technologies) (CyQuant: λ_ex_: 489 ± 12nm / λ_em_: 530 ± 10nm; PrestoBlue: λ_ex_: 547 ± 12nm / λ_em_: 612 ± 20nm). The viability of cells for each drug was normalized to the DMSO vehicle control. In the Figures, results from the CyQuant reading are shown, because they reflect viable cell numbers. Experiments in hypoxic conditions were performed in 1% O_2_ (Heracell 150 incubator, Thermo Fisher Scientific).

The 3D spheroid screens were performed in six technical replicates. Each plate included 7 of the 42 compounds and 6 controls as described for 2D, only the concentration of Encorafenib was increased to 1 µM. The cells were seeded in 96-well ultralow attachment plates (faCellitate) in 150 µl medium and cultured for 4 days to allow for a dense sphere formation. The cell seeding numbers were adjusted to obtain sphere sizes of 370-400 µm after 4 days. This size of spheres has been shown to present different proliferative zones and to have a hypoxic area in its centre (Friedrich et al., 2009). Following 4-day sphere pre-formation, 50% of the medium was replaced by fresh MPM and drugs were added (higher concentration C2 only) for 96h. To allow the drugs to penetrate spheres, we decided to prolong the assay time to 96h. Sphere size and appearance was monitored every day with brightfield imaging on a Cytation 5 (Agilent Technologies). After 96h, the spheres were lysed to perform the CellTiter 3D Glo (Promega, ATP readout) assay according to the manufacturer’s instructions. Luciferase signals were measured on a Cytation 5.

### Determination of IC_50_ values

Cells were seeded in black-walled 96-well µclear plates (Greiner Bio-one) for 2D adherent cell growth. After 3 days drug treatment, cell viability was detected using the CyQuant Direct Cell Proliferation Assay Kit (λ_ex_: 489 ± 12nm / λ_em_: 530 ± 10nm) and PrestoBlue ( λ_ex_: 547 ± 12nm / λ_em_: 612 ± 20nm) with a parallel detection protocol as described above. Blank values (medium only) were subtracted for each well and values were normalized to the untreated control. To determine the respective IC_50_ values, data were plotted and IC_50_ calculated using GraphPad Prism 9, log [inhibitor] vs. response-variable slope 4PL curve fit.

### Seahorse Real-time ATP rate assay

The Seahorse XFe96 analyser (Agilent Technologies) was used to measure oxygen consumption and extracellular acidification rates as described for the Real-time ATP rate assay (Agilent Technologies). Briefly, 10,000 or 20,000 cells for 624Mel and WM3248, respectively, were seeded in a poly-D-lysine-coated Seahorse 96-well plate in RPMI. On the day of the assay, cells were washed with Seahorse XF RPMI (pH 7.4) supplemented with 10 mM glucose, 2 mM glutamine, and 1 mM pyruvate (all Agilent Technologies), and incubated in a non-CO_2_ incubator at 37°C for 1h. Sequential injections of 1.5 µM Oligomycin and 0.5 µM Rotenone/AntimycinA allowed calculation of glycolytic and mitochondrial ATP production rates according to manufacturer’s instructions.

### 2D live/dead cell assay upon drug treatment

Cells were seeded in 96-well flat-bottom plates (Greiner Bio-one). Treatments were applied at concentrations and times indicated in the Figure legends. At the assay endpoint the cells were stained with Hoechst 33342 (1 µg/mL; nuclear live cell stain, Life Technologies) and Sytox Orange (1 µM; nuclear dead cell stain, Life Technologies) for 30min and imaged on the Cytation 5 (Agilent Technologies) using DAPI and RFP filters/LED cubes. The Gen5 software (Agilent Technologies) was used to determine total cell numbers (using Hoechst-stained nuclei) and dead cell numbers (Sytox orange-positive nuclei).

### Determination of the NAD_+_/NADH ratio

10,000 cells per well were seeded in 96-well plates and permitted to adhere. Inhibitors were added and following 24h incubation, NAD^+^ and NADH was quantified using the NAD/NADH-Glo Assay kit (Promega, G9072) according to the manufacturer’s instructions. Luminescence was measured using the Cytation 5 (Agilent Technologies).

### 3D cell viability assays

624Mel-DR-GFP and WM3248-DR-GFP cells were seeded in 96-well ultralow attachment U-bottom plates (FaCellitate) in Mel-MPM for spheroid growth. After two days, half of the culture medium was replaced with fresh medium with or without drugs. To track spheroid growth over time, brightfield images of the spheroids were taken regularly on a Cytation 5. After 4 days of treatment, cells were stained with Propidium Iodide (PI, 4 µg/mL) for 3h to check for cell death within the spheroids. Sphere size, GFP (as a live cell indicator), and PI were measured on the Cytation 5 instrument. Image processing was performed using the Agilent BioTek Gen5 software. GFP mean intensities per pixel were calculated within a plug. Six technical replicates were analysed for each condition.

### Cell culture in OrganoPlates

1x10^4^ WM3248-DR-GFP, 2.5x10^3^ HMEC1-BFP cells and 2.5x10^3^ NHDF-iRFP cells were embedded per µl of Geltrex LDEV-free reduced growth factor basement membrane matrix ( a soluble form of basement membrane extracted from murine Engelbreth-Holm-Swarm tumours; Gibco, final concentration around 9 mg/mL). 2 µl of the cell-matrix mix was added to each gel inlet channel of a 3lane40 OrganoPlate (Mimetas). The gel was solidified for 15min at 37°C and Mel- MPM with 2.5% FBS was added to the medium channels. After 2 days pre-incubation, the medium was replaced with fresh Mel-MPM with or without FK866. The medium was exchanged after 2 days with fresh Mel-MPM with or without FK866. The OrganoPlates were kept on a shaker in the incubator (7° inclination, shaking interval time 8min). Four days after drug addition, the cells were imaged on a Cytation 10 instrument (Agilent Technologies).

## Results

### Generation of drug-resistant melanoma cell lines by prolonged drug exposure

As targeted therapy of BRAF-mutated melanoma with BRAF/MEK-inhibitors inevitably results in drug resistance, we worked with pairs of drug-sensitive and drug-resistant (DR) melanoma cell lines (Figure 1). BRAF-mutant 624Mel, WM3248 and A375 cells were made resistant against the combined treatment of Encorafenib and Binimetinib, a BRAF/MEK-inhibitor combination recently approved by the FDA (Dummer et al., 2018; Shirley, 2018). First, we performed dose-response analyses of the drug-naïve melanoma cells for each compound individually. Representative dose-response curves for the drug-sensitive lines are depicted in Figures 2A (black lines) and the corresponding IC_50_ values in Figure 2B. All cell lines showed low nanomolar IC_50_ values. Melanoma cells under continuous drug exposure (approx. 10 -times the IC_50_ concentrations) initially stopped proliferation and underwent cell line-specific phenotypic alterations (data not shown) before starting to re-proliferate after several weeks / months of combination treatment (Figure 1). Compared to the drug-naïve lines, the drug-resistant ones showed clearly distinct dose-response curves. Cell growth was either only inefficiently suppressed or growth suppression only occurred at much higher concentrations of the respective drug (Figure 2A; red lines). To evaluate if the resistance phenotype was maintained without selective pressure, drugs were withdrawn from the drug-resistant (DR) cells for 3 weeks. Figure 2C shows that the dose-response curves of cells under continuous exposure and of those deprived of drugs for 3 weeks superimpose, indicating that the DR phenotype is stable.

### Drug-resistant cell lines show increased mitochondrial ATP production in comparison to their drug-naïve counterparts

Two pairs of cell lines, 624Mel/624Mel-DR and WM3248/WM3248-DR were selected for a first evaluation of possible metabolic changes during the development of drug resistance. The respective contributions of glycolysis and oxidative phosphorylation to ATP production in drug-sensitive, short-term drug-treated, and drug-resistant cells were assessed by Seahorse ATP production rate assays. Encorafenib/Binimetinib treatment initially led to a downregulation of glycolysis as seen by the drop in glycolytic ATP production after 1 day, which was even more pronounced after 3 days. In contrast, drug-resistant cells resume glycolytic ATP production and show an increased mitochondrial ATP production compared to their sensitive counterparts ( Figure 2D). Thus, our Encorafenib/Binimetinib-resistant cell lines recapitulate the metabolic feature already described for melanoma cells resistant to other BRAF/MEK1 inhibitor combinations (Alkaraki et al., 2021; Avagliano et al., 2020; Pendleton et al., 2023) and encouraged us to perform the screen with a metabolism-oriented compound library.

### The most potent compounds suppress melanoma cell viability at or below concentrations of 1 μM in 2D cell culture

We performed a compound screen with a library of 42 compounds under a number of conditions (RPMI *vs*. MPM medium; hypoxia *vs*. normoxia; 2D *vs*. 3D) using two drug concentrations, C1 and C2 (C2 = 10 x C1), which were individually selected for each drug based on published literature (see Figure S1 and Table S1).

As potency is a desired feature of a successful drug, we first checked how the drugs performed at the lower concentration C1. Figure S2 summarises the viability values obtained in the 2D culture compound screen comparing RPMI (left part) to the more physiologic medium MPM (right part). Figure S2A shows that some drugs had already strong effects at lower concentrations. The NAMPT inhibitor FK866, the HIF inhibitor BAY87-2243 and DHODH inhibitor BAY2402234 had nanomolar activity (50 nM, 100 nM and 50 nM respectively), suppressing cell viability below 50% of controls for most lines (highlighted in bold). Torin2, an mTOR inhibitor, likewise suppressed cell viability by > 50% of controls for all lines at 1 µM (highlighted in bold; Figure S2A).

Other compounds showed cell line-specific effects (highlighted in grey in Figure S2, for further discussion see below): e.g. the Glutaminase inhibitor CB839 (2 µM) efficiently suppressed viability of drug-resistant WM3248 and A375 cells in RPMI while the GPX4 inhibitor ML162 (1 µM) specifically inhibited viability of WM3248 cells in RPMI, independent of their drug resistance status. The dihydrofolate reductase inhibitors Methotrexate and Permetrexed were also very efficient at 1 µM, especially in A375 cells.

As expected, at higher concentrations, more drugs showed strong growth suppression. The GPX4 inhibitor ML162 (10 µM) suppressed the viability of all cell lines in both media. The same was observed for CPI613 (200 µM) and Torin2 (10 µM) (highlighted in bold, Figure S2B). FK866, BAY 87-2243, and BAY2202234 did not result in enhanced effects compared to the 10-fold lower C1 concentration (grey writing). Other interesting compounds were identified, e.g. FX11, BZ-423 and UK5099 (green text in Figure S2B), which seemed to be more effective in MPM, while Erastin showed the inverse behaviour (highlighted in blue; Figure S2B). For a more detailed analysis of these effects, see below.

In conclusion, FK866, BAY87-2243, BAY2402234, as well as Torin2 were the most potent compounds in the 2D screen, acting on the viability of most cell lines and conditions.

### Acquired drug resistance and metabolic rewiring: most compounds performing differently in drug-sensitive and drug-resistant cells do so in a cell line-specific and culture condition-dependent manner

As resistance to therapy presents a major clinical caveat for melanoma patients, we compared the drug response profiles of sensitive and DR cells for each culture condition: 2D in normoxia, in RPMI or MPM; 2D in hypoxia in MPM; 3D in normoxia in MPM (Figure S3, same rows).

The comparison of the drug responses for each cell line under the same environmental conditions (Figure S3) indicates cell type-specific differences between drug-sensitive versus -resistant cells, as illustrated by the differential effects observed for some compounds . For example, in MPM, AZD7545 (PDHK inhibitor) suppressed growth more efficiently in 624MelDR and A375DR than in sensitive cells, while in the WM3248 cell line it was more potent in sensitive cells (Figure S3, second row).

Comparing the drug response for the same pairs of drug-sensitive and -resistant cell lines under different growth conditions (Figure S3, left, middle, or right panels), it becomes apparent that the culture conditions have a strong influence on compound performance. For example, in 624Mel cells, compound C reduces the viability of sensitive cells to a higher extent when cultured in 2D (in RPMI, MPM normoxia and hypoxia) but has a higher potency in DR cells in 3D cultures ( Figure S3, left panels). For WM3248 cells, BAY-2402234 (DHODH inhibitor) reduces viability more efficiently in DR cells in 2D cultures in MPM (normoxia and hypoxia), but this effect is not visible under 3D culture conditions or in RPMI medium ( Figure S3, middle panels).

The variability of the results underscores the importance of environmental factors, such as nutrient levels, oxygen concentrations and 3D growth, when screening metabolism-targeting compounds. In conclusion, the acquired drug resistance clearly affected the cellular sensitivities to several drugs. However, these effects were not consistently observed for all three pairs of cell lines and not under all culture conditions. On the other hand, some drugs potently suppressed growth of both, sensitive and DR cells (see below). Therefore, we subsequently focused on the differential effects of drugs observed between the distinct culture conditions.

### Identification of compounds with differential activity in physiological MPM vs. standard RPMI

Figure 3A depicts the results of all six cell lines in 2D, highlighting those compounds which show a differential efficacy in RPMI or in MPM while Figure 3B presents the ratio, RPMI/MPM, of the % viability for all 42 compounds, with higher activity in MPM indicated by brighter yellow colour (see also Figure S4 and S5 for more results in RPMI *vs*. MPM).

**Figure 3.**
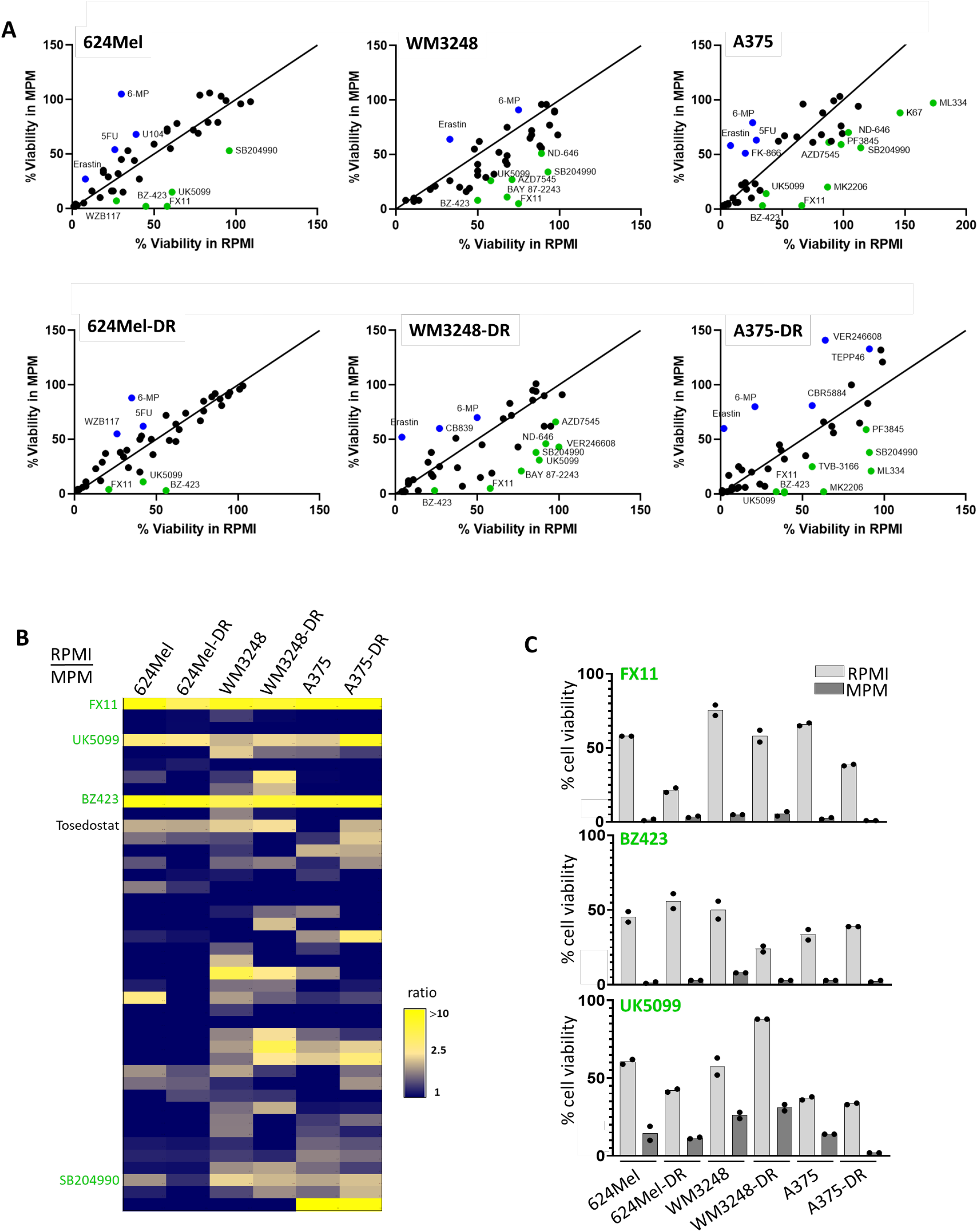
**Compounds showing different efficacies in RPMI vs. MPM**. A) Plots of % viability (from the 2D screen) in RPMI *vs*. MPM for each cell line. Compounds highlighted in blue are more potent in RPMI, compounds highlighted in green are more potent in MPM. The threshold for highlighting the compounds was ≥ 50% viability reduction in MPM or RPMI, with a minimum of 20% viability difference between both. The data plotted correspond to the mean of the technical replicates of the screen for the higher concentration C2. B) Heatmap representation of the ratio of the % viability RPMI/MPM (see Figure S4 for further details). Yellow colour indicates a higher ratio and thus more efficacy of the compound in MPM. The compounds showing a ratio ≥ 2 in at least 4 lines are indicated and those above the threshold described in A are highlighted in green. C) Bar diagram representing cell viabilities from the 2D screen (concentration C2) for all six lines.

FX-11 (reported to be an inhibitor of LDHA), BZ-423 (an ATPase inhibitor), UK5099 (an inhibitor of the mitochondrial pyruvate carrier (MPC1) show a considerably higher activity in the physiological medium, MPM, across all cell lines (Figure 3A-C).

Other drugs generally had higher activity in RPMI (Figure 3A (blue dots), Figure S4B, Figure S5B), such as Erastin (targeting the cysteine-glutamate transporter SLC7A11 (xCT)), FK866 (a nicotinamide phosphoribosyltransferase (NAMPT) inhibitor), the glutaminase inhibitor (GLS) CB- 839 (in WM3248DR and A375DR cells), and the pyrimidine analogue 5FU.

### Serum albumin binding is responsible for the lower efficacy of FX-11 and BZ-423 in RPMI

Next, we further investigated the cause of the differential activity of FX-11 and BZ-423, two compounds showing stronger effects in MPM than in RPMI. Both compounds are of interest as they are reported to affect two major metabolic pathways: FX-11, described to target LDH and therefore glycolysis, and BZ-423, described to inhibit ATP synthase and thereby OXPHOS. First, we validated the effects observed in the screen by delineating dose -response curves of both compounds in sensitive and DR 624Mel and WM3248 cells, in RPMI and MPM, in biological triplicates. Representative results are shown in Figure S6A, confirming that FX-11 and BZ-423 inhibit growth more efficiently in MPM, as opposed to RPMI, across a wide range of concentrations. Major differences between MPM and RPMI are the glutamine concentration (higher in RPMI) and the fact that MPM was designed to be a xeno -reduced medium with only 2.5% FCS instead of 10% for RPMI (Preis et al., 2024). To check if any of the above-mentioned differences were responsible for this effect, we supplemented MPM so that the glutamine concentration was 2 mM and FCS concentration was 10%. Furthermore, MPM (containing 2.5% FCS) was supplemented with albumin to approximately match the albumin concentration present in 10% FCS (Figure S6B). It has been previously described that melanoma cells rely on glutaminolysis in commercial medium (Baenke et al., 2016) and the glutamine concentration is 4 times higher in RPMI than in MPM (Preis et al., 2024). However, increasing the glutamine concentration in MPM to RPMI levels did not shift the dose response curves. On the contrary, the addition of 10% FBS to MPM rendered the dose response curves very similar to the ones obtained for RPMI. Finally, albumin addition to the medium had the same effect ( Figure S6B and S6C). These data indicate that FX-11 and BZ-423 likely bind to albumin. Although both drugs are interesting as they inhibit cell growth in all culture conditions, FX-11 was not investigated further here, since we found FX-11 to have an alternative function compared to its published target, lactate dehydrogenase (data not shown). Since BZ-423 was also only active at concentrations between 10 and 20 µM, we focussed on other more potent compounds for further validations.

### Oxygen concentration influences compound efficacy in MPM

Hypoxia is common in the core of solid tumours, due to the large distance to vessels and disrupted tumour vasculature. Reduced oxygen concentrations have multiple effects on cancer cells and their tumour microenvironment: hypoxia alters intercellular communication via exosomes (Walbrecq et al., 2020) and has been directly linked to drug resistance in melanoma (Almeida et al., 2019; Qin et al., 2016) presenting a major obstacle to ameliorate patient outcomes (Codony et al., 2021). Comparing the results from the screen under normoxic (MPM) and hypoxic conditions (MPM-1% O_2_), we observed that most compounds show reduced efficacy under hypoxia; generally, cells seem to be more resistant in oxygen-reduced conditions (Figure S7A). The growth inhibitory effect e.g. of the HIF-1 inhibitor BAY87-2243 was consistently reduced under hypoxia when HIF-1 levels were increased (Figures 4B and S7). In addition to the HIF-1 inhibitor, also inhibitors of other protein targets known to be upregulated by hypoxia showed reduced potency. These include, for example, the glucose transporter (SLC2A1,3,4) inhibitor WZB117, the pyruvate dehydrogenase kinase (PDHK) inhibitor AZD7545, and the NRF2 inhibitor ML385. In addition to BAY87-2243, the potency of the two other compounds with nanomolar activity, BAY2402234 (DHODH inhibitor) and FK866 (NAMPT inhibitor) ( Figure 4B and 4C) was also reduced under hypoxic conditions in 2D, clearly indicating the importance to routinely assess efficacies of drugs targeting solid tumour growth also under oxygen-reduced conditions.

**Figure 4.**
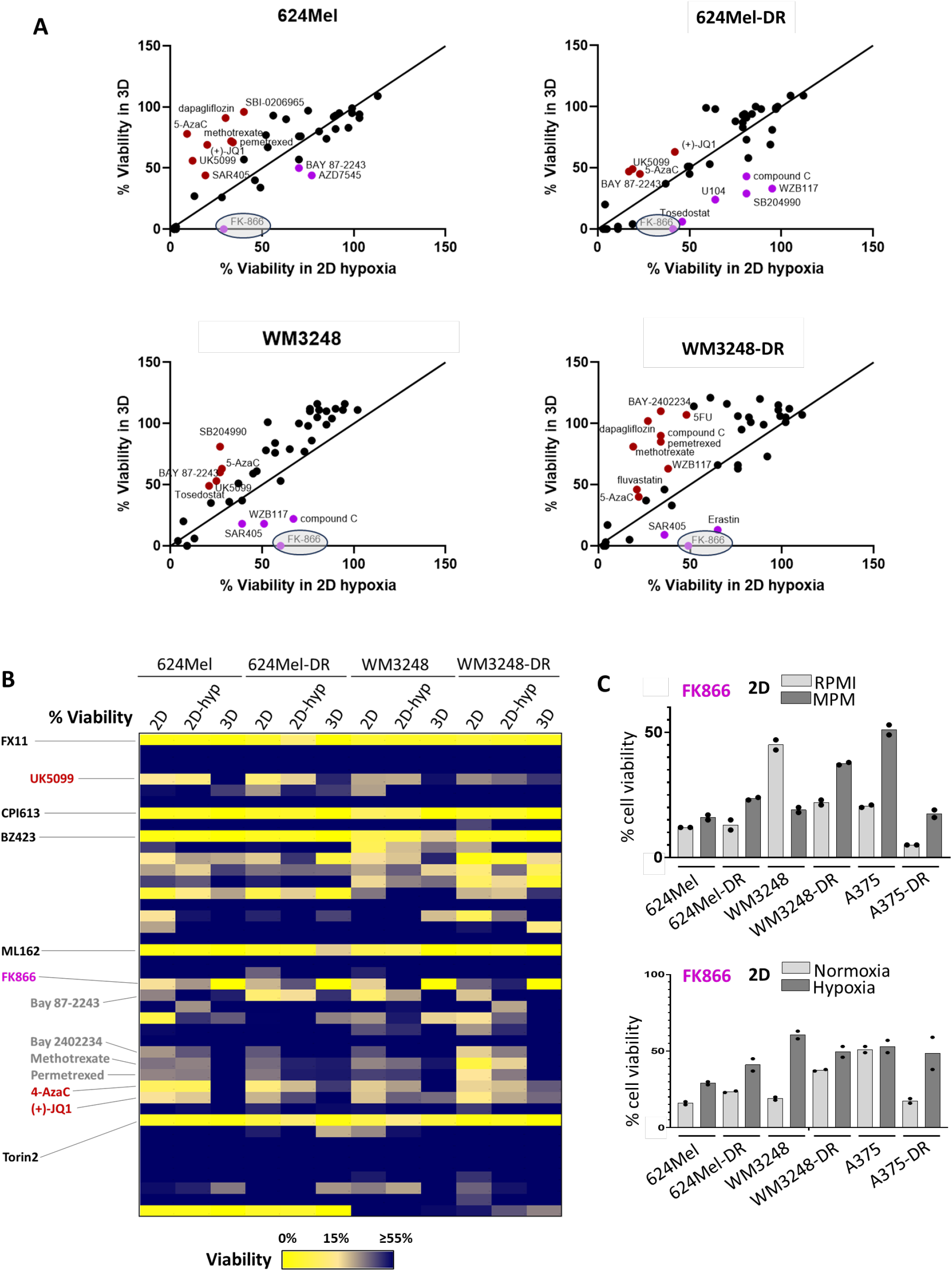
The NAMPT inhibitor FK866 has a higher efficacy in 3D spheroid assays compared to its effects in 2D under normoxia or hypoxia. A) Plots of % viability values (from the screen in MPM) comparing 2D-hypoxia vs. 3D spheroids for each cell line. Compounds highlighted in blue are more potent in 3D spheroids, compounds highlighted in dark red are more potent in hypoxic 2D cultures. Threshold for highlighting the compounds were ≥ 50% viability reduction in 2D (hypoxia) or 3D, with a minimum of 20 % viability difference between both. The data plotted correspond to the mean of the technical replicates of the screen for the higher concentration C2. B) Heatmap representation of the % viability for the indicated conditions. Yellow colour indicates a lower viability compared to untreated and thus more efficacy of the compound in this condition. The compounds above the threshold described in A are highlighted in blue or dark red text. Other compounds discussed in the text are highlighted in black or grey text. C) Bar diagrams representing cell viability for the NAMPT inhibitor, FK866, in the 2D screen comparing RPMI and MPM (top panel) and in 2D in MPM under normoxia and hypoxia (concentration C2) (lower panel).

### The compounds Torin 2 and FK866 show the highest efficacy in suppressing growth in 3D conditions

Against a trend of reduced activity of many compounds in 3D spheroids, 6 compounds retained high efficacy, one of which, FK866, was even more potent in 3D compared to 2D conditions. Drug effects were compared for the three screening conditions in physiological medium: (i) MPM- 2D, (ii) MPM-2D-1% O_2,_ and (iii) MPM-3D. We hypothesised that such comparisons might inform about compounds performing best in more complex 3D systems. Overall, many drugs performed less efficiently in melanoma spheroids compared to monolayer cultures ( Figure 4A and B). Drugs that performed well in monolayer cultures but not in 3D included UK5099 (MPC1i), 5-AzaC (DNA methylation inhibitor) and (+)-JQ1 (bromodomain inhibitor) (highlighted in red in Figure 4A and B). Interestingly, also many of the most potent compounds showing nanomolar activity in 2D (see also Figure S2A) did not efficiently suppress 3D spheroid growth, e.g. BAY87-2243, BAY2402234, Methotrexate and Permetrexed (highlighted in grey in Figure 4B).

However, we also identified five compounds which were still effective in all physiological conditions including 3D spheroids: FX-11 (described to target LDH), CPI613 (targeting α- ketoglutarate dehydrogenase (αKGDH) and pyruvate dehydrogenase (PDH)), BZ -423 (ATPase inhibitor), ML-162 (Glutathione peroxidase 4 (GPX4) inhibitor) and Torin2 (mTOR inhibitor) (black annotation in Figure 4B), of which Torin2 was the only compound to suppress viability at low micromolar concentrations (C1) in the 2D screen (see Figure S2A).

Most interestingly, the NAMPT inhibitor FK866 (highlighted in blue in Figure 4), was even more efficient in suppressing growth of 3D melanoma spheroids compared to 2D conditions. In 2D assays, FK866 showed nanomolar activity, was generally more effective in RPMI compared to MPM and was generally less effective under hypoxia compared to normoxia (see Figure 4C). Of note, for FK866 the 2D hypoxia condition does not reflect the behaviour in 3D. Due to their potency in 2D cultures and 3D spheroids, Torin2 and FK866 were further characterised (see below).

### Combining Torin2 with an autophagy inhibitor does not show decreased cell viability in 3D spheroid assays

Torin2, an mTOR inhibitor which was very potently inhibiting both 2D and 3D growth of our melanoma lines, was further characterised. Since inhibition of mTOR induces autophagy, which can rescue cancer cells under stress, we hypothesised that additional inhibition of autophagy might synergise with Torin2 to suppress melanoma cell viability (Figure 5A). We first tested the effect of 3 autophagy inhibitors (SAR405, SBI0206965 and MRT68921) individually on our drug resistant cells in 2D viability assays using MPM as culture medium. SAR405, a VSP35 inhibitor, and SBI0206965, an ULK1/2 inhibitor had considerable effects on cell viability in the screen (e.g. Figure S2B, lane 12 and 13). MRT68921, targeting ULK1/2, was the most potent compound and suppressed viability efficiently below 10 µM (Figure 5B) as single treatment.

**Figure 5.**
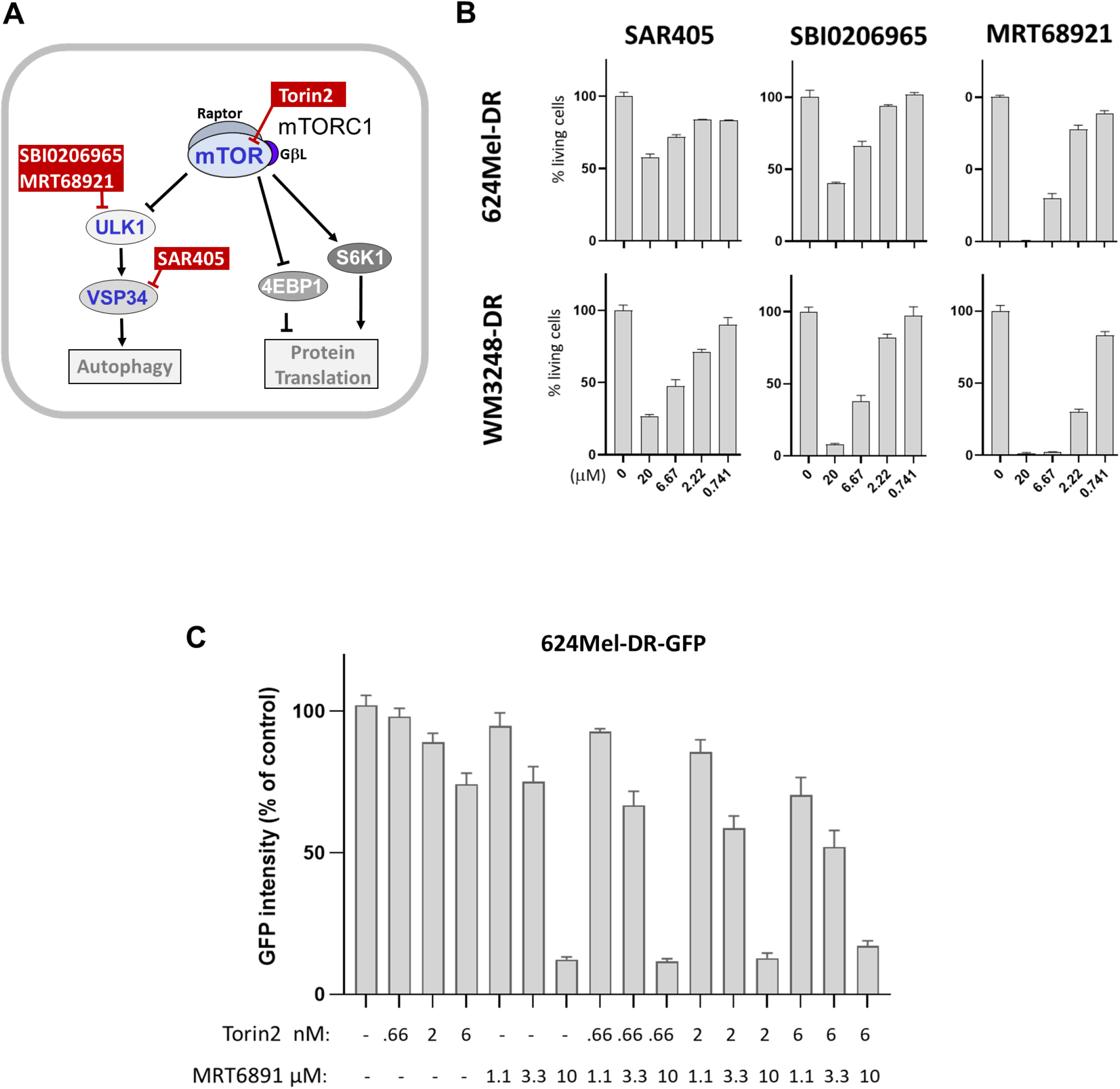
Targeting melanoma cells by Torin2, alone and in combination with autophagy inhibitors. A) Scheme showing the proteins and pathways targeted by the co-treatment strategy. B) Effect of 3 different autophagy inhibitors on the viability of 2D cultures of WM3248-DR and 624Mel-DR cells. Viability was assessed using the Hoechst33342/Sytox Orange assay and was represented as % of untreated control. Mean and standard deviation of one representative biological replicate out of 3 are shown, each performed in 3 technical replicates. C) Effects of single treatments and co-treatment of Torin2 and MRT68921 on 3D spheroid cultures of 624Mel- DR-GFP cells. Mean GFP intensity is represented as % of untreated control. Mean and standard deviation of one representative biological replicate out of 3 are shown, each performed in 6 technical replicates.

We then investigated the viability of 3D spheroid cultures by exposing them to different Torin2 or MRT68921 concentrations. Torin2 reduced spheroid size efficiently at nanomolar concentrations (Figure 5C). We exposed the 624Mel-DR-GFP 3D spheroids to three low Torin2 concentrations at which the spheroid size began to be affected by Torin2 treatment (0.66 to 6 nM) and to a range of MRT68921 concentrations. Unexpectedly, we did not see decreased cell viability compared to the single treatments when we combined both treatments (Figure 5C).

### The NAMPT inhibitor FK866 very potently inhibits the viability of drug-resistant melanoma cells also in a matrix-embedded multi-cell-type 3D system

The NAMPT inhibitor FK866 was one of the most interesting compounds identified from the screen as it showed nanomolar activity in the 2D (Figure S3A) and the 3D spheroids screen (Figure 4). NAMPT is a crucial enzyme in the NAD salvage pathway, on which cancer cells especially rely on (Yaku et al., 2018), and its inhibition leads to a reduction of all NAD metabolites. To test the on-target effect of FK866, we measured NAD^+^ and NADH levels in untreated *vs*. FK866-treated WM3248 and 624Mel cells and demonstrated that FK866 reduced the levels of both metabolites drastically (Figure 6A).

**Figure 6.**
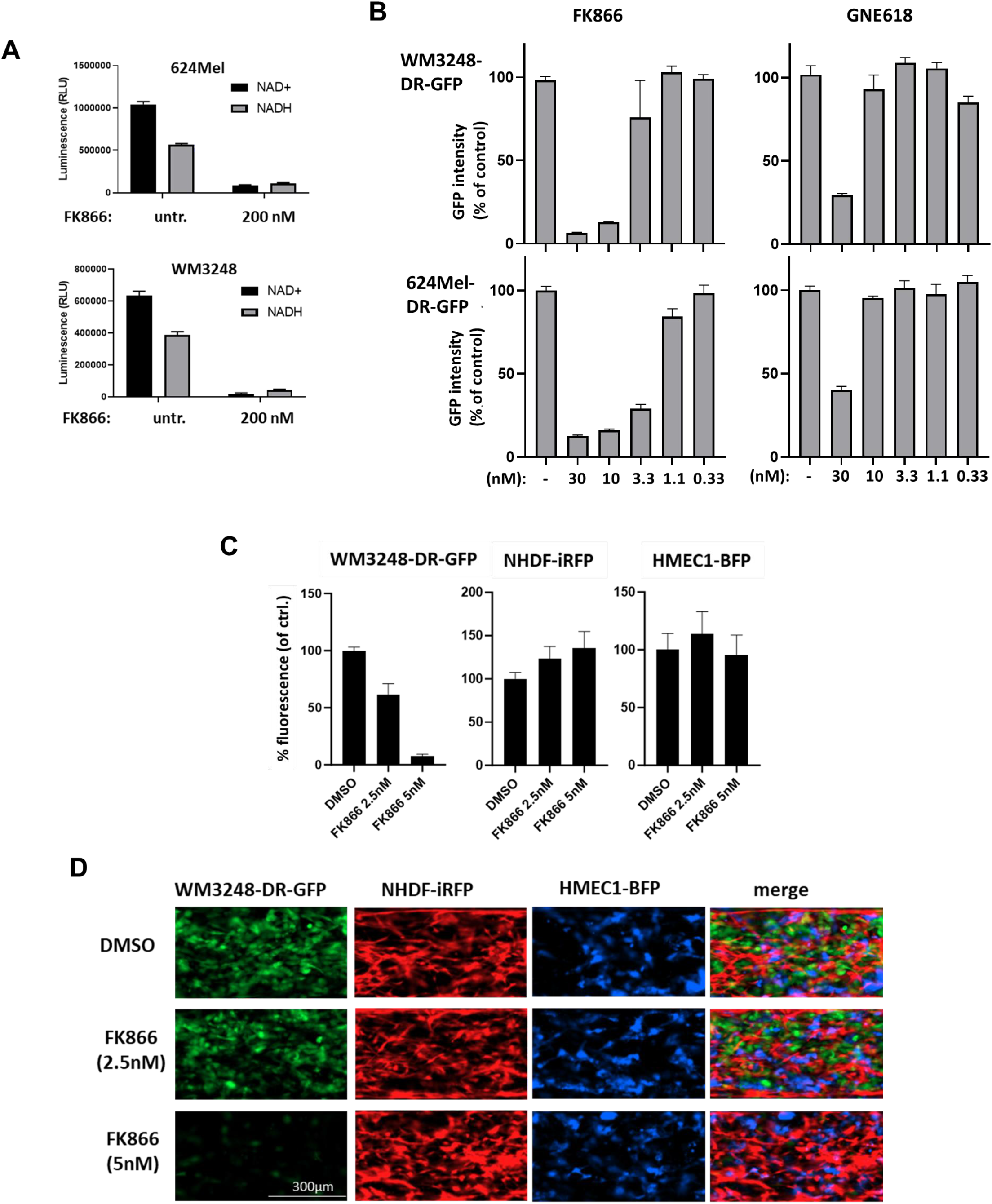
The NAMPT inhibitor FK866 very potently and specifically inhibits the growth of melanoma cells. A) Effects of 200 nM of FK866 on the levels of NAD+ and NADH using the NAD/NADH-Glo Assay kit (Promega, G9072). Mean and standard deviation of one representative biological replicate out of 3 are shown, each performed in 4 technical replicates. B) Effects of various FK866 and GNE618 concentrations on 3D spheroid cultures of 624Mel-DR-GFP and WM3248-DR-GFP. Mean GFP intensity is represented as % of untreated control. Mean and standard deviation of one representative biological replicate out of 3 are shown (except for GNE619: n=2), each performed with 6 technical replicates. C) Effect of FK866 on co-cultures of WM3248-DR-GFP, HMEC1-BFP and NHDF-iRFP cells embedded in Geltrex matrix in an 3lane40 OrganoPlate (Mimetas). The data are normalised to the DMSO treated control for each cell line. Mean and standard deviation of one representative biological replicate out of 3 are shown, each performed in 4 technical replicates. D) Images corresponding to the OrganoPlate assay described in C.

In 3D spheroid assays we included two different NAMPT inhibitors, FK866 and GNE618, and showed that FK866 more potently supressed 3D spheroid viability than GNE618 (Figure 6B). 624Mel-DR-GFP-spheroids were slightly more sensitive to FK866 treatment compared to WM3248-DR-GFP-spheroids (Figure 6B).

To test if FK866 treatments can also potently suppress melanoma growth in culture systems which more closely reflect the tumour environment, we used amatrix-embedded model combining melanoma cells, fibroblasts and endothelial cells in OrganoPlates^®^ (Mimetas) (as described before (Preis et al., 2024)). Briefly, we used GFP-labelled WM3248-DR, iRFP-labelled normal human dermal fibroblasts (NHDF-iRFP), and TagBFP-labelled human microvascular endothelial cells (HMEC1-BFP), which were embedded in Geltrex matrix (a soluble form of basement membrane extracted f rom murine Engelbreth-Holm-Swarm tumours), seeded into the microchannels of an OrganoPlate and incubated with the physiologic medium (Mel-MPM). The different cell types can be visualized according to their specific fluorophores (expressed when the cells are viable) to assess the effects of the drug treatments. FK866 reduced WM3248 -DR-GFP viability quite drastically at a concentration of 5 nM, while not influencing the viability of the labelled NHDF -iRFP or HMEC1-BFP cells (Figure 6C and D). Thus, this observation suggests that there might be a therapeutic window for the use of NAMPT inhibitors in drug-resistant melanoma, since we could target our most resilient cell line WM3248-DR specifically while not affecting healthy bystander cells.

## Discussion

Genetic mutations and nutrient availability determine the growth behaviour of cancer cells (reviewed in Kanarek et al., 2020; Sullivan & Vander Heiden, 2019; Vander Heiden & DeBerardinis, 2017). Cells of a given tissue type harbouring specific mutations are therefore vulnerable to drugs targeting specific metabolic pathways. In BRAF -mutated melanoma treatment, the emergence of resistance occurring after targeted therapy (combined BRAF and MEK inhibition) has been associated with the occurrence of metabolic rewiring (Alkaraki et al., 2021; Avagliano et al., 2020).

Using BRAF-mutated melanoma cells we generated BRAFi/MEKi ( Encorafenib/Binimetinib) resistant cells, which showed increased mitochondrial ATP production in comparison to their drug-naïve counterparts, thereby recapitulating ametabolic adaptation known to occur in drug-resistant melanoma. We subjected the drug-sensitive and -resistant cell lines to a screen with a metabolically targeted compound library using a self -made physiologic medium (MPM) which we compared to RPMI. We recently described that compounds eliciting ferroptosis show a different potency in MPM *vs*. RPMI (Preis et al., 2024) and thus expected some differences in the drug screening, also since nutrient concentration differences between standard media and physiologic media would be expected to influence metabolic pathways. Along these lines, differential effects of drugs have recently been described for drug efficacies in standard cell culture media *vs*. physiologic media or sera (Abbott et al., 2023; Biancur et al., 2017; Cantor et al., 2017; Muir et al., 2017) and in blood versus lymph (Ubellacker et al., 2020). Interestingly, a study describing a large compound screen on 482 cell lines cultured either in RPMI or a modified adult bovine serum (ftABS) demonstrated that drugs targeting metabolic proteins displayed differential potency in RPMI compared to ftABS, while compounds targeting signalling proteins did not ( Abbott et al., 2023).

Our screen was designed to dissect differences in compound activity comparing the following conditions (for drug-sensitive and drug-resistant cells): (i) RPMI *vs*. MPM medium in 2D cultures (normoxia only) to identify drugs which act differently in the two media (different nutrient levels / FBS concentration), (ii) normoxia *vs*. hypoxia in 2D cultures (MPM only) to identify drugs changing efficacy with low O_2_, a condition characteristic of solid tumours, (iii) 2D cultures (normoxia and hypoxia) *vs*. 3D using the MPM medium to identify drugs which are influenced by nutrient/oxygen gradients and/or perform less well due to decreased penetrance into the denser structure of spheroids mimicking the architecture of solid tumours. From these screening data we then selected the most potent drugs for further validation in 2D, 3D spheroids or a more complex multi-cell type 3D system.

Concerning differences of drug behaviour that can be attributed to the different cell culture media, we were able to identify compounds with higher efficacy in standard RPMI medium. For example, A375-DR and WM3248-DR cells were more sensitive to the glutaminase inhibitor CB-839 at both concentrations used (2 and 20 µM) when cultured in RPMI (Figure S2A and S5C) than in MPM. This result is in line with previous publications showing stronger glutamine addiction of resistant melanoma cells in standard medium (Baenke et al., 2016). Glutaminase inhibitors have also been reported in several preclinical studies to target different types of cancer cells *in vitro,* but to a lesser extent *in vivo* (Biancur et al., 2017). In addition, glutaminase inhibitors were reported to perform poorly *in vitro* when cells were cultured in serum-like medium or in a modified adult bovine serum (Abbott et al., 2023; Muir et al., 2017). Moreover, for three of the six cell lines (624Mel, 624Mel-DR, A375), we observed that 5-FU was less potent in MPM (Figure S5C), an observation already reported before for acute myeloid leukaemia cells cultured in HPLM medium in comparison to RPMI (Cantor et al., 2017).

Besides the recapitulation of specific previously described effects of drugs in standard *vs.* physiologic media we also identified additional inhibitors with distinct responses in the different media. Higher compound efficacy in standard RPMI medium was found for the xCT inhibitor Erastin, the purine analogue 6-MP (Figure 3A, S4 and S5B), and the NAMPT inhibitor FK-866 (Figure 4C). On the other hand, we also identified drugs whose efficacy is greater in MPM, e.g. the LDH inhibitor FX11, the mitochondrial F _1_F_0-_ATPase inhibitor BZ-423, the mitochondrial pyruvate carrier (MPC) inhibitor UK5099 (Figure 3), the alanyl aminopeptidase (ANPEP) inhibitor Tosedostat and the ATP citrate lyase (ACLY) inhibitor SB204990 (Figure 3A and B and S5A). The IC_50_’s of FX-11 and BZ-423 in our melanoma cell lines were consistently lower in MPM ( Figure S6A). However, since the screen in MPM was performed under lower levels of FCS (2.5% *vs.* 10% in RPMI), we suspected that drugs binding plasma proteins might show up in this assay format in this condition. Supplementing FCS or albumin to MPM to match the concentration contained in 10% FCS had the effect of raising the IC_50_ observed in MPM to the one in RPMI (Figure S6B and C). Our motivation to reduce the FBS levels in MPM to 2.5% was mainly to reduce the animal burden by using a xeno-reduced, more physiological medium, since our cells showed a similar growth rate in MPM with 2.5% FBS compared to RPMI with 10% FBS. However, the discrepancies observed for FX11 and BZ-423, due to different amounts of FBS in both media, highlights the importance of using the same serum concentrations for reliable comparisons of different media. It also highlights a general weakness of *in vitro* screens, no matter in which medium they are performed. Many drugs potentially bind serum proteins and as reviewed before (Liu et al., 2014; Smith et al., 2010), this is not a reason to exclude those compounds from further validation steps, since the free drug concentration *in vivo* rather depends on drug clearance mechanisms (mostly involving the liver) than on plasma protein binding. In *in vitro* systems, however, the free drug concentration is dictated by the concentration of plasma proteins (Liu et al., 2014; Smith et al., 2010). Thus, we suggest performing screens in the presence of a lower and a higher amount of FBS/albumin to detect albumin binding compounds, which have to be processed differently during validations.

As outlined before, there is a dire need to identify 2^nd^ line treatment options for patients who have become resistant to first line drugs. However, comparing drug efficacies in drug-sensitive and - resistant cells, we found that most differences were cell line-specific and depended on culture conditions (Figure S3), so that drugs inhibiting all resistant cell lines were not identified.

Not surprisingly, many compounds were less effective under hypoxia, especially those compounds for which the target protein has been described to be upregulated by hypoxia, e.g. the HIF-1 inhibitor BAY87-2243, the glucose transporter (SLC2A1,3,4) inhibitor WZB117 and the pyruvate dehydrogenase kinase (PDHK) inhibitor AZD7545 (Figure S7).

Overall, many compounds also showed reduced efficacy in melanoma spheroids compared to monolayer cultures (Figure 4A and B), e.g. UK5099 (MPC1i), 5-AzaC (DNA methylation inhibitor) and (+)-JQ1 (bromodomain inhibitor), BAY87-2243, BAY2402234, Methotrexate and Permetrexed. Also, these results were not unexpected as they likely reflect the effects of hypoxic conditions within the spheroid combined with impaired penetrance of certain chemical scaffolds into the dense structure of the spheroid.

Together, these findings indicate that many drugs have reduced efficacy in conditions which more closely resemble the *in vivo* situation, again stressing the importance to improve pre-clinical drug screening and validation systems to better mimic physiological conditions found in patients.

As potency is a desired feature of a successful drug, we focussed on identifying drugs which perform at concentrations below 1 µM in 2D cell cultures. The most potent compounds were the NAMPT inhibitor FK866, the HIF inhibitor BAY87-2243, the DHODH inhibitor BAY2402234, which had nanomolar activity (50 nM, 100 nM and 50 nM, respectively), as well as the mTOR inhibitor Torin2 (1 µM) (Figure S2A). Of those very potent compounds, Torin2 was effective in all physiological conditions including 3D spheroids (Figure 4B) and the NAMPT inhibitor, FK866 (Figure 4), was even more efficient to suppress growth in 3D spheroids compared to 2D conditions. Thus, due to their potency in 2D cultures and 3D spheroids , Torin2 and FK866 were further characterised.

The inhibition of mTOR can induce autophagy, which can enable cells to cope with stress (Maes & Agostinis, 2014). To test whether a combination of mTOR and autophagy inhibition would further decrease cell viability, we treated cells in 3D with Torin2 and MRT68921. Unfortunately, this drug combination did not perform better than the single treatments (Figure 5C).

The NAMPT inhibitor, FK866, performed better in RPMI compared to MPM, which is in line with the recent findings by others (Abbott et al., 2023), describing reduced efficacy of FK866 in serum-based media. In MPM under hypoxia, FK866 was also less effective compared to normoxic conditions, while the compound was very effective in 3D spheroid assays. This shows that applying hypoxia to 2D cultures is too simple an approach to recapitulate the behaviour of compounds in 3D spheroids. We could show that FK866 treatment greatly reduces the amount of NAD+ and NADH in our cell lines (Figure 6A), suggesting that the compound inhibits its proposed target. In summary, FK866 potently suppressed 2D and 3D cell viability in drug-resistant melanoma cell lines at nanomolar concentrations. A co-culture assay of matrix embedded drug resistant WM3248-DR-GFP melanoma cells (which was the most resistant cell line to FK866 in 2D and 3D spheroid cultures), with dermal fibroblasts (NHDF-iRFP) and endothelial cells (HMEC1-BFP), showed that at concentrations as low as 5 nM, FK866 drastically suppressed melanoma cell growth, while having no effect on the growth of fibroblasts or endothelial cells (Figure 6C and D). This suggests that there may be a therapeutic window to target the drug resistant melanoma cells over other cell types in the tumour. NAMPT has emerged as interesting drug target in a number of tumours (Nomura et al, 2023, Nature; Galli et al, 2020, Frontiers). NAMPT inhibitors have been proposed to be more toxic to cancer cells than to healthy tissue since the NAD *de novo* synthesis is not enough to cope with the cancer cells’ high demand of this metabolite. Therefore, tumour cells are more reliant on the NAMPT-dependent salvage pathway. Still, until today, no NAMPT inhibitor has been approved for clinical use due to toxicity and poor response rates (Holen et al., 2008; Von Heideman et al., 2010) in clinical trials. However, more recent advances in developing an antibody-coupled NAMPT inhibitor, that is targeted to the tumour site, has shown promising results *in vivo* (Böhnke et al., 2022). Specific targeting to melanoma cell expressed antigens such as MCSP (also called chondroitin sulfate proteoglycan 4 (CSPG4)), a transmembrane cell-surface molecule, which is highly expressed in most human melanomas (Campoli et al., 2010; Pitcovski et al., 2017; Price et al., 2011) could enhance the therapeutic window *in vivo*. Based on our results, NAMPT inhibitors specifically targeting melanoma cells represent promising and new targeted therapy options for melanoma patients who do not respond to current therapies or have become resistant.

## Acknowledgements

The 624Mel cells were a kind gift of Dr. Ruth Halaban (Dermatology Department, Yale School of Medicine, USA) and WM35 were gifted from Dr. Markus Böhm (University of Münster, Germany). We thank Dr. Fabrice Tolle for viral transduction of melanoma cells with GFP-luc and Dr. Martin Nurmik for help with the sorting of the GFP expressing melanoma lines. We thank Dr. Joanna Wroblewska for viral transduction and sorting of NHDF and HMEC cells.

This work was supported by the Luxembourg National Research Fund (FNR), project number 10675146 (funding scheme PRIDE, “CANBIO”), and by the Fondation Cancer Luxembourg (“SecMelPro” grant).

## References

1. Abbott, K. L., Ali, A., Casalena, D., Do, B. T., Ferreira, R., Cheah, J. H., Soule, C. K., Deik, A., Kunchok, T., Schmidt, D. R., Renner, S., Honeder, S. E., Wu, M., Chan, S. H., Tseyang, T., Greaves, D., Ng, C. W., Clish, C. B., Rees, M. G., … Muir, A. (2023). Screening in serum-derived medium reveals differential response to compounds targeting metabolism. BioRxiv. 10.1101/2023.02.25.529972

2. Abbott, K. L., Ali, A., Casalena, D., Do, B. T., Ferreira, R., Cheah, J. H., Soule, C. K., Deik, A., Kunchok, T., Schmidt, D. R., Renner, S., Honeder, S. E., Wu, M., Chan, S. H., Tseyang, T., Stoltzfus, A. T., Michel, S. L. J., Greaves, D., Hsu, P. P., … Vander Heiden, M. G. (2023). Screening in serum-derived medium reveals differential response to compounds targeting metabolism. Cell Chemical Biology, 30(9). 10.1016/j.chembiol.2023.08.007

3. Alkaraki, A., McArthur, G. A., Sheppard, K. E., & Smith, L. K. (2021). Metabolic plasticity in melanoma progression and response to oncogene targeted therapies. Cancers, 13(22). 10.3390/cancers13225810

4. Almeida, F. V., Douglass, S. M., Fane, M. E., & Weeraratna, A. T. (2019). Bad company: Microenvironmentally mediated resistance to targeted therapy in melanoma. In Pigment Cell and Melanoma Research (Vol. 32, Issue 2). 10.1111/pcmr.12736

5. Ascierto, P. A., Casula, M., Bulgarelli, J., Pisano, M., Piccinini, C., Piccin, L., Cossu, A., Mandalà, M., Ferrucci, P. F., Guidoboni, M., Rutkowski, P., Ferraresi, V., Arance, A., Guida, M., Maiello, E., Gogas, H., Richtig, E., Fierro, M. T., Lebbe, C., … Palmieri, G. (2024). Sequential immunotherapy and targeted therapy for metastatic BRAF V600 mutated melanoma: 4 -year survival and biomarkers evaluation from the phase II SECOMBIT trial. Nature Communications, 15(1). 10.1038/s41467-023-44475-6

6. Avagliano, A., Fiume, G., Pelagalli, A., Sanità, G., Ruocco, M. R., Montagnani, S., & Arcucci, A. (2020). Metabolic Plasticity of Melanoma Cells and Their Crosstalk With Tumor Microenvironment. Frontiers in Oncology, 10(May), 1–21. 10.3389/fonc.2020.00722

7. Baenke, F., Chaneton, B., Smith, M., Van Den Broek, N., Hogan, K., Tang, H., Viros, A., Martin, M., Galbraith, L., Girotti, M. R., Dhomen, N., Gottlieb, E., & Marais, R. (2016). Resistance to BRAF inhibitors induces glutamine dependency in melanoma cells. Molecular Oncology, 10(1), 73–84. 10.1016/j.molonc.2015.08.003

8. Biancur, D. E., Paulo, J. A., Małachowska, B., Del Rey, M. Q., Sousa, C. M., Wang, X., Sohn, A. S. W., Chu, G. C., Gygi, S. P., Harper, J. W., Fendler, W., Mancias, J. D., & Kimmelman, A. C. (2017a). Compensatory metabolic networks in pancreatic cancers upon perturbation of glutamine metabolism. Nature Communications, 8. 10.1038/ncomms15965

9. Biancur, D. E., Paulo, J. A., Małachowska, B., Del Rey, M. Q., Sousa, C. M., Wang, X., Sohn, A. S. W., Chu, G. C., Gygi, S. P., Harper, J. W., Fendler, W., Mancias, J. D., & Kimmelman, A. C. (2017b). Compensatory metabolic networks in pancreatic cancers upon perturbation of glutamine metabolism. Nature Communications, 8(May). 10.1038/ncomms15965

10. Böhnke, N., Berger, M., Griebenow, N., Rottmann, A., Erkelenz, M., Hammer, S., Berndt, S., Günther, J., Wengner, A. M., Stelte-Ludwig, B., Mahlert, C., Greven, S., Dietz, L., Jörißen, H., Barak, N., Bömer, U., Hillig, R. C., Eberspaecher, U., Weiske, J., … Sommer, A. (2022). A Novel NAMPT Inhibitor-Based Antibody-Drug Conjugate Payload Class for Cancer Therapy. Bioconjugate Chemistry, 33(6), 1210–1221. 10.1021/acs.bioconjchem.2c00178

11. Campoli, M., Ferrone, S., & Wang, X. (2010). Functional and Clinical Relevance of Chondroitin Sulfate Proteoglycan 4. In Advances in Cancer Research (Vol. 109, Issue C). 10.1016/B978-0-12-380890-5.00003-X

12. Cantor, J. R., Abu-Remaileh, M., Kanarek, N., Freinkman, E., Gao, X., Louissaint, A., Lewis, C. A., & Sabatini, D. M. (2017a). Physiologic Medium Rewires Cellular Metabolism and Reveals Uric Acid as an Endogenous Inhibitor of UMP Synthase. Cell, 169(2), 258–272.e17. 10.1016/j.cell.2017.03.023

13. Colombino, M., Capone, M., Lissia, A., Cossu, A., Rubino, C., De Giorgi, V., Massi, D., Fonsatti, E., Staibano, S., Nappi, O., Pagani, E., Casula, M., Manca, A., Sini, M. C., Franco, R., Botti, G., Caracò, C., Mozzillo, N., Ascierto, P. A., & Palmieri, G. (2012). BRAF/NRAS mutation frequencies among primary tumors and metastases in patients with melanoma. Journal of Clinical Oncology, 30(20). 10.1200/JCO.2011.41.2452

14. Czarnecka, A. M., Bartnik, E., Fiedorowicz, M., & Rutkowski, P. (2020). Targeted therapy in melanoma and mechanisms of resistance. In International Journal of Molecular Sciences (Vol. 21, Issue 13). 10.3390/ijms21134576

15. Dummer, R., Ascierto, P. A., Gogas, H. J., Arance, A., Mandala, M., Liszkay, G., Garbe, C., Schadendorf, D., Krajsova, I., Gutzmer, R., Chiarion-Sileni, V., Dutriaux, C., de Groot, J. W. B., Yamazaki, N., Loquai, C., Moutouh-de Parseval, L. A., Pickard, M. D., Sandor, V., Robert, C., & Flaherty, K. T. (2018). Encorafenib plus binimetinib versus vemurafenib or encorafenib in patients with BRAF-mutant melanoma (COLUMBUS): a multicentre, open-label, randomised phase 3 trial. The Lancet Oncology, 19(5), 603–615. 10.1016/S1470-2045(18)30142-6

16. Friedrich, J., Seidel, C., Ebner, R., & Kunz-Schughart, L. A. (2009). Spheroid-based drug screen: Considerations and practical approach. Nature Protocols, 4(3), 309–324. 10.1038/nprot.2008.226

17. Holen, K., Saltz, L. B., Hollywood, E., Burk, K., & Hanauske, A. R. (2008). The pharmacokinetics, toxicities, and biologic effects of FK866, a nicotinamide adenine dinucleotide biosynthesis inhibitor. Investigational New Drugs, 26(1), 45–51. 10.1007/s10637-007-9083-2

18. Kanarek, N., Petrova, B., & Sabatini, D. M. (2020). Dietary modifications for enhanced cancer therapy. In Nature (Vol. 579, Issue 7800). 10.1038/s41586 -020-2124-0

19. Kozar, I., Margue, C., Rothengatter, S., Haan, C., & Kreis, S. (2019). Many ways to resistance : How melanoma cells evade targeted therapies. BBA - Reviews on Cancer, 1871(2), 313–322. 10.1016/j.bbcan.2019.02.002

20. Lee, S. H., Hu, W., Matulay, J. T., Silva, M. V., Owczarek, T. B., Kim, K., Chua, C. W., Barlow, L. M. J., Kandoth, C., Williams, A. B., Bergren, S. K., Pietzak, E. J., Anderson, C. B., Benson, M. C., Coleman, J. A., Taylor, B. S., Abate-Shen, C., McKiernan, J. M., Al-Ahmadie, H., … Shen, M. M. (2018). Tumor Evolution and Drug Response in Patient-Derived Organoid Models of Bladder Cancer. Cell, 173(2), 515–528.e17. 10.1016/j.cell.2018.03.017

21. Lelliott, E. J., McArthur, G. A., Oliaro, J., & Sheppard, K. E. (2021). Immunomodulatory Effects of BRAF, MEK, and CDK4/6 Inhibitors: Implications for Combining Targeted Therapy and Immune Checkpoint Blockade for the Treatment of Melanoma. In Frontiers in Immunology (Vol. 12). 10.3389/fimmu.2021.661737

22. Liu, X., Wright, M., & Hop, C. E. C. A. (2014). Rational use of plasma protein and tissue binding data in drug design. Journal of Medicinal Chemistry, 57(20). 10.1021/jm5007935

23. Maes, H., & Agostinis, P. (2014). Autophagy and mitophagy interplay in melanoma progression. Mitochondrion, 19(Part A), 58–68. 10.1016/j.mito.2014.07.003

24. Muir, A., Danai, L. V., Gui, D. Y., Waingarten, C. Y., Lewis, C. A., & Vander Heiden, M. G. (2017a). Environmental cystine drives glutamine anaplerosis and sensitizes cancer cells to glutaminase inhibition. ELife, 6, 1–27. 10.7554/eLife.27713

25. Muir, A., Danai, L. V., & Vander Heiden, M. G. (2018). Microenvironmental regulation of cancer cell metabolism: Implications for experimental design and translational studies. In DMM Disease Models and Mechanisms (Vol. 11, Issue 8). 10.1242/dmm.035758

26. Pendleton, K. E., Wang, K., & Echeverria, G. V. (2023). Rewiring of mitochondrial metabolism in therapy-resistant cancers: permanent and plastic adaptations. In Frontiers in Cell and Developmental Biology (Vol. 11). 10.3389/fcell.2023.1254313

27. Pitcovski, J., Shahar, E., Aizenshtein, E., & Gorodetsky, R. (2017). Melanoma antigens and related immunological markers. In Critical Reviews in Oncology/Hematology (Vol. 115). 10.1016/j.critrevonc.2017.05.001

28. Preis, J. R., Rolvering, C., Kirchmeyer, M., Halilovic, D., Wroblewska, J. P., Tolle, F., Behrmann, I., & Haan, C. (2024). Ferroptosis susceptibility of melanoma cells: dependence on cell-type, acquired drug resistance, and medium composition. BioRxiv. 10.1101/2024.08.12.607529

29. Price, M. A., Colvin Wanshura, L. E., Yang, J., Carlson, J., Xiang, B., Li, G., Ferrone, S., Dudek, A. Z., Turley, E. A., & McCarthy, J. B. (2011). CSPG4, a potential therapeutic target, facilitates malignant progression of melanoma. In Pigment Cell and Melanoma Research (Vol. 24, Issue 6). 10.1111/j.1755-148X.2011.00929.x

30. Qin, Y., Roszik, J., Chattopadhyay, C., Hashimoto, Y., Liu, C., Cooper, Z. A., Wargo, J. A., Hwu, P., Ekmekcioglu, S., & Grimm, E. A. (2016). Hypoxia-driven mechanism of vemurafenib resistance in melanoma. Molecular Cancer Therapeutics, 15(10). 10.1158/1535-7163.MCT-15-0963

31. Shirley, M. (2018). Encorafenib and Binimetinib: First Global Approvals. Drugs, 78(12), 1277– 1284. 10.1007/s40265-018-0963-x

32. Smith, D. A., Di, L., & Kerns, E. H. (2010). The effect of plasma protein binding on in vivo efficacy: Misconceptions in drug discovery. In Nature Reviews Drug Discovery (Vol. 9, Issue 12). 10.1038/nrd3287

33. Sullivan, M. R., & Vander Heiden, M. G. (2019). Determinants of nutrient limitation in cancer. In Critical Reviews in Biochemistry and Molecular Biology (Vol. 54, Issue 3). 10.1080/10409238.2019.1611733

34. Ubellacker, J. M., Tasdogan, A., Ramesh, V., Shen, B., Mitchell, E. C., Martin-Sandoval, M. S., Gu, Z., McCormick, M. L., Durham, A. B., Spitz, D. R., Zhao, Z., Mathews, T. P., & Morrison, S. J. (2020). Lymph protects metastasizing melanoma cells from ferroptosis. Nature, 585(7823). 10.1038/s41586-020-2623-z

35. Vande Voorde, J., Ackermann, T., Pfetzer, N., Sumpton, D., Mackay, G., Kalna, G., Nixon, C., Blyth, K., Gottlieb, E., & Tardito, S. (2019). Improving the metabolic fidelity of cancer models with a physiological cell culture medium. Science Advances, 5(1), eaau7314. 10.1126/sciadv.aau7314

36. Vander Heiden, M. G., & DeBerardinis, R. J. (2017). Understanding the Intersections between Metabolism and Cancer Biology. In Cell (Vol. 168, Issue 4). 10.1016/j.cell.2016.12.039

37. Vlachogiannis, G., Hedayat, S., Vatsiou, A., Jamin, Y., Fernández-Mateos, J., Khan, K., Lampis, A., Eason, K., Huntingford, I., Burke, R., Rata, M., Koh, D., Tunariu, N., Collins, D., Hulkki - Wilson, S., Ragulan, C., Spiteri, I., Moorcraft, S. Y., Chau, I., … Valeri, N. (2018). Patient-derived organoids model treatment response of metastatic gastrointestinal cancers. Science, 359(6378), 920–926. 10.1126/science.aao2774

38. Von Heideman, A., Berglund, Å., Larsson, R., Nygren, P., & Larsson, R. (2010). Safety and efficacy of NAD depleting cancer drugs: Results of a phase i clinical trial of CHS 828 and overview of published data. Cancer Chemotherapy and Pharmacology, 65(6), 1165–1172. 10.1007/s00280-009-1125-3

39. Walbrecq, G., Margue, C., Behrmann, I., & Kreis, S. (2020). Distinct cargos of small extracellular vesicles derived from hypoxic cells and their effect on cancer cells. International Journal of Molecular Sciences, 21(14). 10.3390/ijms21145071

40. Whiteman, D. C., Green, A. C., & Olsen, C. M. (2016). The Growing Burden of Invasive Melanoma: Projections of Incidence Rates and Numbers of New Cases in Six Susceptible Populations through 2031. Journal of Investigative Dermatology, 136(6), 1161–1171. 10.1016/j.jid.2016.01.035

41. Winder, M., & Virós, A. (2018). Mechanisms of drug resistance in Melanoma. In Handbook of Experimental Pharmacology (Vol. 249). 10.1007/164_2017_17

42. Yaku, K., Okabe, K., Hikosaka, K., & Nakagawa, T. (2018). NAD metabolism in cancer therapeutics. Frontiers in Oncology, 8(DEC), 1–9. 10.3389/fonc.2018.00622

