## Supplemental table and figure legends for "Metabolism-oriented compound screen in physiological culture conditions identifies a NAMPT inhibitor highly effective against drug-naïve and -resistant melanoma cells"

**Table S1. Information about the compounds and concentrations, C1 and C2, used in the screen.**

**Figure S1. Pie chart showing major metabolic pathways targeted by the library compounds.**

**Figure S2.** **Results of the 2D screen under normoxia.** Compounds were used A) at the lower concentration C1 and B) at the 10-fold higher concentration C2. Yellow colour indicates a lower viability with respect to the untreated control and thus more efficacy of the compound in this condition. Compounds highlighted in black writing are potent in most cell lines and conditions. Compounds indicated in grey show activity in less conditions or show cell line or condition specificity. Compounds highlighted in green perform better in MPM, while the compound in blue performs better in RPMI.

**Figure S3. Comparison of drug responses in sensitive and DR cells**. Plots of % viability values from the screen, comparing sensitive (S) *vs*. DR cells for each cell line and culture condition. Compounds highlighted in red are more potent in DR cells, compounds highlighted in blue are more potent in sensitive cells. The threshold for highlighting the compounds was ≥ 50% viability reduction in S or DR, with a minimum of 20% viability difference between both. The data plotted correspond to the mean of the technical replicates of the screen for the higher concentration C2.

**Figure S4**: **Comparison of drug responses in 2D viability assays using RPMI *vs*. MPM** A) Heatmap representation of the ratio of the % viability RPMI/MPM. Yellow colour indicates a higher ratio and thus more efficacy of the compound in MPM. The names of compounds showing a ratio ≥2 in at least 4 cell lines are annotated in black and those above the threshold described in Figure 3A are highlighted in green. B) Heatmap representation of the ratio of the % viability MPM/RPMI. Yellow colour indicates a higher ratio and thus more efficacy of the compound in RPMI. The names of compounds showing a ratio ≥2 in at least 4 lines are indicated, those above the threshold described in Figure 3A are highlighted in blue.

**Figure S5. Selected compounds with different efficacy in RPMI and MPM.** Bar diagram representation of % cell viability values from the 2D screen for all cell lines. Error bars indicate mean ± SD of two technical duplicates used in the screen. A) Selected compounds from the screen showing a greater efficacy in MPM (concentration C2) (see also Figure 3). B) Selected compounds showing a higher efficacy in RPMI (concentration C2). C) The glutaminase inhibitor CB-893 (concentration C1) shows increased efficacy in specific drug-resistant lines, and the anti-metabolite 5-FU (concentration C2) showed increased efficacy in RPMI in certain lines.

**Figure S6. The serum albumin concentration influences the efficacy of FX-11 and BZ-423**. Viability was assessed using the CyQuant assay and represented as % of untreated. A) Representative dose-response curves of FX-11 and BZ-423 for sensitive and drug-resistant 624Mel and WM3248 cells in RPMI *vs.* MPM. B) Representative dose-response curves of FX-11 and BZ-423 for 624Mel cells in RPMI and MPM with supplements as indicated. C) IC_50_ values derived from biological replicates of FX-11- or BZ-423-treated 624Mel cells. The values were derived from the curves using Graphpad Prism 9. 2 to 5 biological replicates were carried out, each in 3 technical replicates.

**Figure S7. Comparison of drug effects in normoxia and hypoxia.**

A) Plots of % viability values from the screen, comparing cells grown in 2D in MPM under normoxia to those cultivated under hypoxia. Compounds highlighted in red are more potent in normoxia, compounds highlighted in dark red are more potent in hypoxia. The threshold for highlighting the compounds was ≥ 50% viability reduction in normoxia or hypoxia, with a minimum of 20% viability difference between both. The data plotted correspond to the mean of the technical replicates of the screen for the higher concentration C2. B) Selected compounds from the screen showing reduced potency when melanoma cells were cultured under hypoxia. Viability was assessed using the Cyquant assay and the viability is represented as % of untreated.
