## Supplemental tables and figures for "Metabolism-oriented compound screen in physiological culture conditions identifies a NAMPT inhibitor highly effective against drug-naïve and -resistant melanoma cells"

### Table S1

##### Concentrations of compounds used for the screen

| Pos. | compound | Molecular target | action | category | C1 (μM) | C2 (μM) |
|---|---|---|---|---|---|---|
| 1 | FX11 | LDHA | inhibitor | Glycolysis | 2.5 | 25 |
| 2 | TEPP46 | PKM2 | activator | Glycolysis | 2.5 | 25 |
| 3 | TLN232 | PKM2 | inhibitor | Glycolysis | 5 | 50 |
| 4 | UK5099 | SLC54A1 and 2 | inhibitor | TCA | 10 | 100 |
| 5 | AZD7545 | PDHK | inhibitor | TCA | 2 | 20 |
| 6 | CB839 | GLS | inhibitor | TCA | 2 | 20 |
| 7 | CPI613 | PDH and αKGDH/OGDH | inhibitor | TCA | 20 | 200 |
| 8 | VER246608 | PDHK | inhibitor | TCA | 1 | 10 |
| 9 | BZ-423 | ATP synthase | inhibitor | Oxidative Phosphorylation | 2 | 20 |
| 10 | CBR5884 | PHGDH | inhibitor | Amino acid met. | 5 | 50 |
| 11 | Tosedostat | ANPEP and LAP3 | inhibitor | Amino acid met. | 1 | 10 |
| 12 | SAR405 | VSP34 | inhibitor | autophagy | 2 | 20 |
| 13 | SBI-0206965 | Ulk1 /2 | | autophagy | 2 | 20 |
| 14 | fluvastatin | HMGCR | inhibitor | Mevalonate pathway | 1 | 10 |
| 15 | A-769662 | AMPK | activator | AMPK | 2 | 20 |
| 16 | compound C | AMPK | inhibitor | AMPK | 0.75 | 7.5 |
| 17 | Erastin | SLC7A11 | inhibitor | Cystin import | 1.5 | 15 |
| 18 | K67 | p62/SQSTM | inhibitor | oxidative stress | 1.5 | 15 |
| 19 | ML-162 | GPX4 | inhibitor | oxidative stress | 1 | 10 |
| 20 | ML334 | NRF2 | activator | oxidative stress | 2 | 20 |
| 21 | ML385 | NRF2 | inhibitor | oxidative stress | 2 | 20 |
| 22 | FK-866 | NAMPT | inhibitor | NAD metabolism | 0.05 | 0.5 |
| 23 | BAY 87-2243 | HIF | inhibitor | HIF1 | 0.1 | 1 |
| 24 | dapagliflozin | SLC5A2/1 | inhibitor | Glycolysis | 2 | 20 |
| 25 | WZB117 | SLC2A4/1/3 | inhibitor | Glycolysis | 5 | 50 |
| 26 | 5FU | TYMS | inhibitor | 1C met./ nucleotide met. | 4 | 40 |
| 27 | 6MP | PPAT | inhibitor | 1C met./ nucleotide met. | 5 | 50 |
| 28 | BAY-2402234 | DHODH | inhibitor | 1C met./ nucleotide met. | 0.05 | 0.5 |
| 29 | methotrexate | DHFR | inhibitor | 1C met./ nucleotide met. | 1 | 10 |
| 30 | pemetrexed | GARFT / DHFR/ TS | inhibitor | 1C met./ nucleotide met. | 1 | 10 |
| 31 | 5-AzaC | DNA methylation | inhibitor | DNA methylation | 1 | 10 |
| 32 | (+)-JQ1 | BRD | inhibitor | c-myc inhibition | 1 | 10 |
| 33 | epacadostat | IDO1 | inhibitor | Amino acid met. | 5 | 50 |
| 34 | Torin2 | mTor | inhibitor | Akt-mTOR | 1 | 10 |
| 35 | U104 | CA | inhibitor | pH homeostasis | 10 | 100 |
| 36 | CP 597396 | SLC9A1 | inhibitor | pH homeostasis | 2 | 20 |
| 37 | JZL184 | MGLL/MAGL | inhibitor | fatty acid met. | 1 | 10 |
| 38 | PF3845 | FAAH | inhibitor | fatty acid met. | 1 | 10 |
| 39 | ND-646 | ACC | inhibitor | fatty acid met. | 1 | 10 |
| 40 | SB204990 | ACLY | inhibitor | fatty acid met. | 5 | 50 |
| 41 | TVB-3166 | FASN | inhibitor | fatty acid met. | 1 | 10 |
| 42 | MK2206 | Akt | inhibitor | Akt-mTOR | 1 | 10 |

### Figure S1

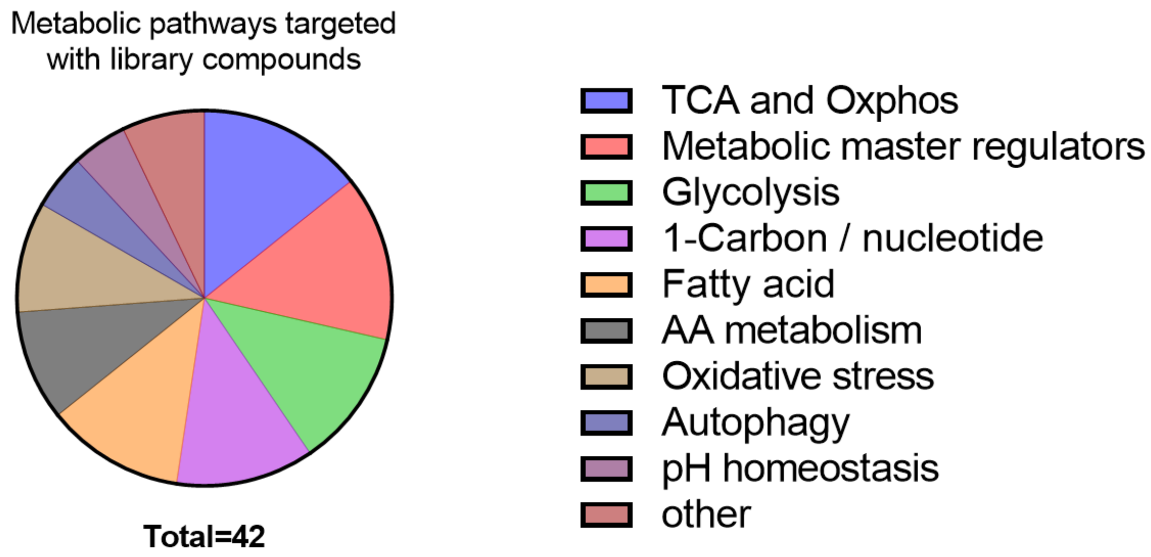
Metabolic pathways targeted
with library compounds
Total=42
TCA and Oxphos
Metabolic master regulators
Glycolysis
1-Carbon / nucleotide
Fatty acid
AA metabolism
Oxidative stress
Autophagy
pH homeostasis
other

#### Figure S2

### A

C1: Low concentration

| Annotation | % viability | RPMI 624Mel Sens. | RPMI 624Mel DR | RPMI Wm3248 Sens. | RPMI Wm3248 DR | RPMI A375 Sens. | RPMI A375 DR | MPM 624Mel Sens. | MPM 624Mel DR | MPM Wm3248 Sens. | MPM Wm3248 DR | MPM A375 Sens. | MPM A375 DR | # |
|---|---|---|---|---|---|---|---|---|---|---|---|---|---|---|
| | FX11 | 103 | 91 | 106 | 89 | 95 | 93 | 103 | 90 | 97 | 99 | 84 | 70 | 1 |
| | TEPP46) | 98 | 94 | 101 | 103 | 101 | 99 | 112 | 93 | 96 | 96 | 82 | 94 | 2 |
| | TLN232 | 106 | 95 | 107 | 99 | 96 | 100 | 113 | 101 | 95 | 97 | 89 | 102 | 3 |
| | UK5099 | 94 | 71 | 101 | 107 | 74 | 68 | 84 | 63 | 99 | 90 | 81 | 67 | 4 |
| | AZD7545 | 83 | 84 | 109 | 115 | 86 | 78 | 78 | 72 | 88 | 93 | 85 | 84 | 5 |
| CB839 (GLS) | CB839 | 85 | 65 | 97 | 9 | 70 | 6 | 84 | 69 | 94 | 86 | 70 | 39 | 6 |
| | CPI613 | 101 | 99 | 106 | 75 | 102 | 95 | 114 | 93 | 70 | 88 | 90 | 100 | 7 |
| | VER246608 | 108 | 105 | 99 | 97 | 100 | 97 | 103 | 103 | 87 | 103 | 99 | 101 | 8 |
| | BZ-423 | 95 | 107 | 105 | 110 | 95 | 94 | 103 | 103 | 84 | 104 | 87 | 93 | 9 |
| | CBR5884 | 91 | 81 | 24 | 49 | 86 | 66 | 105 | 91 | 37 | 75 | 99 | 63 | 10 |
| | Tosedostat | 81 | 69 | 72 | 110 | 21 | 45 | 66 | 49 | 83 | 99 | 12 | 35 | 11 |
| | SAR405 | 64 | 59 | 84 | 51 | 71 | 89 | 33 | 46 | 81 | 65 | 51 | 59 | 12 |
| | SBI-0206965 | 93 | 99 | 92 | 78 | 90 | 93 | 106 | 92 | 110 | 89 | 76 | 100 | 13 |
| | fluvastatin | 83 | 33 | 97 | 83 | 40 | 86 | 85 | 15 | 94 | 100 | 27 | 100 | 14 |
| | A-769662 | 100 | 106 | 108 | 118 | 101 | 98 | 98 | 107 | 101 | 106 | 98 | 100 | 15 |
| | compound C | 103 | 103 | 103 | 115 | 99 | 99 | 87 | 102 | 101 | 100 | 107 | 110 | 16 |
| | Erastin | 75 | 93 | 86 | 27 | 22 | 75 | 101 | 99 | 92 | 102 | 88 | 100 | 17 |
| | K67 | 98 | 107 | 114 | 119 | 105 | 98 | 101 | 105 | 95 | 104 | 96 | 108 | 18 |
| ML162 (GPX4) | ML-162 | 94 | 108 | 8 | 4 | 50 | 80 | 80 | 103 | 100 | 105 | 83 | 114 | 19 |
| | ML334 | 107 | 105 | 113 | 101 | 111 | 101 | 108 | 104 | 97 | 99 | 107 | 95 | 20 |
| | ML385 | 65 | 75 | 113 | 111 | 67 | 60 | 66 | 51 | 55 | 81 | 71 | 54 | 21 |
| **FK866** (NAMPT) | FK-866 | 24 | 19 | 43 | 26 | 15 | 4 | 17 | 31 | 20 | 50 | 24 | 11 | 22 |
| **Bay 87-2243** (HIF) | BAY 87-2243 | 27 | 8 | 66 | 56 | 12 | 12 | 24 | 10 | 12 | 19 | 14 | 10 | 23 |
| | dapagliflozin | 125 | 99 | 107 | 107 | 101 | 102 | 119 | 108 | 101 | 103 | 101 | 104 | 24 |
| | WZB117 | 119 | 105 | 99 | 106 | 100 | 100 | 113 | 98 | 89 | 104 | 93 | 98 | 25 |
| | 5FU | 68 | 63 | 75 | 58 | 76 | 36 | 117 | 103 | 75 | 100 | 104 | 109 | 26 |
| | 6-MP | 43 | 49 | 66 | 63 | 22 | 13 | 123 | 77 | 94 | 94 | 71 | 60 | 27 |
| **Bay 2402234** (DHODH) | BAY-2402234 | 31 | 34 | 58 | 61 | 6 | 9 | 37 | 39 | 29 | 22 | 6 | 7 | 28 |
| Methotrexate (DHFR) | methotrexate | 28 | 33 | 51 | 36 | 5 | 9 | 32 | 38 | 37 | 10 | 3 | 2 | 29 |
| Permetrexed (DHFR, GARFT, TS) | pemetrexed | 32 | 33 | 57 | 53 | 18 | 25 | 33 | 37 | 51 | 21 | 5 | 3 | 30 |
| | 5-AzaC | 104 | 66 | 82 | 87 | 52 | 45 | 87 | 44 | 78 | 73 | 47 | 28 | 31 |
| | (+)-JQ1 | 61 | 63 | 34 | 26 | 28 | 41 | 69 | 59 | 39 | 42 | 31 | 37 | 32 |
| | epacadostat | 125 | 96 | 107 | 109 | 104 | 104 | 106 | 99 | 98 | 105 | 103 | 113 | 33 |
| **Torin 2** (mTOR) | Torin2 | 4 | 17 | 27 | 9 | 11 | 17 | 4 | 15 | 42 | 25 | 13 | 17 | 34 |
| | U104 | 105 | 101 | 100 | 107 | 99 | 102 | 106 | 93 | 87 | 104 | 94 | 128 | 35 |
| | Zoniporide DHC | 116 | 109 | 108 | 94 | 105 | 103 | 113 | 103 | 89 | 100 | 96 | 106 | 36 |
| | JZL184 | 119 | 110 | 109 | 90 | 108 | 107 | 116 | 102 | 95 | 100 | 94 | 114 | 37 |
| | PF3845 | 126 | 108 | 111 | 112 | 105 | 104 | 124 | 101 | 94 | 101 | 97 | 112 | 38 |
| | ND-646 | 119 | 95 | 98 | 88 | 89 | 55 | 84 | 72 | 69 | 58 | 56 | 35 | 39 |
| | SB204990 | 124 | 111 | 115 | 117 | 106 | 109 | 119 | 103 | 98 | 103 | 112 | 128 | 40 |
| | TVB-3166 | 107 | 87 | 101 | 118 | 84 | 66 | 83 | 69 | 84 | 62 | 61 | 34 | 41 |
| | MK2206 | 108 | 114 | 85 | 72 | 100 | 121 | 85 | 95 | 103 | 112 | 121 | 120 | 42 |

### B

C2: High concentration

| Annotation | % viability | RPMI 624Mel Sens. | RPMI 624Mel DR | RPMI Wm3248 Sens. | RPMI Wm3248 DR | RPMI A375 Sens. | RPMI A375 DR | MPM 624Mel Sens. | MPM 624Mel DR | MPM Wm3248 Sens. | MPM Wm3248 DR | MPM A375 Sens. | MPM A375 DR | # |
|---|---|---|---|---|---|---|---|---|---|---|---|---|---|---|
| **FX11** (LDH) | FX11 | 58 | 21 | 75 | 58 | 66 | 39 | 2 | 4 | 5 | 5 | 3 | 1 | 1 |
| | TEPP46) | 84 | 89 | 94 | 91 | 83 | 91 | 106 | 87 | 77 | 90 | 88 | 133 | 2 |
| | TLN232 | 94 | 95 | 92 | 86 | 92 | 98 | 99 | 93 | 96 | 101 | 99 | 132 | 3 |
| **UK5099** (MPC1/2) | UK5099 | 61 | 42 | 58 | 88 | 37 | 34 | 15 | 11 | 26 | 31 | 14 | 2 | 4 |
| | AZD7545 | 34 | 17 | 71 | 98 | 88 | 39 | 53 | 29 | 27 | 66 | 61 | 32 | 5 |
| | CB839 | 74 | 64 | 68 | 27 | 47 | 11 | 72 | 59 | 75 | 60 | 62 | 25 | 6 |
| **CPI613** (PDH, aKGDH) | CPI613 | 3 | 3 | 11 | 14 | 4 | 1 | 3 | 4 | 10 | 3 | 4 | 3 | 7 |
| | VER246608 | 78 | 84 | 99 | 100 | 67 | 64 | 104 | 89 | 68 | 43 | 96 | 141 | 8 |
| **BZ423** (ATP synth.) | BZ-423 | 45 | 56 | 50 | 24 | 34 | 39 | 2 | 3 | 8 | 3 | 3 | 2 | 9 |
| | CBR5884 | 30 | 41 | 14 | 27 | 52 | 56 | 45 | 53 | 8 | 25 | 67 | 81 | 10 |
| | Tosedostat | 31 | 40 | 43 | 6 | 3 | 6 | 15 | 20 | 16 | 2 | 3 | 3 | 11 |
| | SAR405 | 41 | 48 | 33 | 8 | 28 | 24 | 27 | 36 | 26 | 12 | 23 | 9 | 12 |
| | SBI-0206965 | 38 | 47 | 21 | 8 | 14 | 10 | 44 | 50 | 18 | 10 | 6 | 5 | 13 |
| | fluvastatin | 7 | 7 | 50 | 38 | 6 | 7 | 5 | 7 | 35 | 24 | 5 | 4 | 14 |
| | A-769662 | 91 | 103 | 97 | 84 | 112 | 99 | 103 | 99 | 90 | 95 | 94 | 121 | 15 |
| | compound C | 25 | 58 | 51 | 8 | 17 | 36 | 16 | 49 | 62 | 9 | 22 | 45 | 16 |
| Erastin (xCT) | Erastin | 8 | 68 | 33 | 4 | 8 | 2 | 27 | 74 | 64 | 52 | 58 | 60 | 17 |
| | K67 | 64 | 86 | 97 | 91 | 146 | 80 | 78 | 92 | 89 | 62 | 88 | 100 | 18 |
| **ML162** (GPX4) | ML-162 | 1 | 2 | 6 | 5 | 2 | 1 | 2 | 4 | 8 | 2 | 3 | 1 | 19 |
| | ML334 | 109 | 101 | 89 | 86 | 173 | 92 | 98 | 96 | 96 | 94 | 97 | 21 | 20 |
| | ML385 | 58 | 28 | 68 | 66 | 82 | 37 | 73 | 38 | 49 | 69 | 68 | 40 | 21 |
| FK866 (NAMPT) | FK-866 | 12 | 13 | 45 | 22 | 20 | 5 | 16 | 23 | 19 | 38 | 51 | 17 | 22 |
| Bay 87-2243 (HIF) | BAY 87-2243 | 19 | 7 | 68 | 77 | 32 | 19 | 33 | 11 | 11 | 21 | 17 | 20 | 23 |
| | dapagliflozin | 60 | 79 | 88 | 95 | 97 | 90 | 55 | 85 | 58 | 62 | 103 | 83 | 24 |
| | WZB117 | 27 | 26 | 55 | 23 | 21 | 29 | 7 | 55 | 29 | 16 | 18 | 23 | 25 |
| | 5FU | 26 | 42 | 50 | 20 | 29 | 13 | 54 | 62 | 41 | 29 | 63 | 22 | 26 |
| | 6-MP | 30 | 35 | 75 | 50 | 26 | 21 | 105 | 88 | 91 | 70 | 79 | 80 | 27 |
| Bay 2402234 (DHODH) | BAY-2402234 | 22 | 32 | 68 | 59 | 6 | 7 | 29 | 40 | 41 | 19 | 4 | 6 | 28 |
| | methotrexate | 19 | 19 | 60 | 41 | 12 | 15 | 35 | 37 | 32 | 7 | 6 | 6 | 29 |
| | pemetrexed | 28 | 30 | 67 | 52 | 25 | 27 | 32 | 34 | 44 | 15 | 10 | 7 | 30 |
| | 5-AzaC | 17 | 16 | 39 | 22 | 10 | 4 | 10 | 12 | 20 | 18 | 10 | 2 | 31 |
| | (+)-JQ1 | 24 | 33 | 24 | 11 | 20 | 11 | 16 | 24 | 21 | 13 | 24 | 6 | 32 |
| | epacadostat | 58 | 62 | 63 | 53 | 62 | 63 | 71 | 48 | 52 | 45 | 66 | 66 | 33 |
| **Torin 2** (mTOR) | Torin2 | 2 | 5 | 11 | 4 | 4 | 2 | 4 | 6 | 7 | 2 | 4 | 2 | 34 |
| | U104 | 39 | 40 | 50 | 71 | 75 | 52 | 68 | 50 | 31 | 72 | 61 | 35 | 35 |
| | Zoniporide DHC | 50 | 62 | 89 | 84 | 98 | 68 | 59 | 64 | 56 | 86 | 76 | 62 | 36 |
| | JZL184 | 103 | 94 | 83 | 70 | 99 | 85 | 96 | 90 | 72 | 83 | 69 | 65 | 37 |
| | PF3845 | 89 | 90 | 83 | 102 | 98 | 89 | 79 | 81 | 68 | 91 | 59 | 59 | 38 |
| | ND-646 | 83 | 79 | 89 | 92 | 104 | 69 | 79 | 76 | 51 | 46 | 70 | 56 | 39 |
| | SB204990 | 96 | 77 | 93 | 86 | 114 | 91 | 53 | 67 | 34 | 38 | 56 | 38 | 40 |
| | TVB-3166 | 77 | 56 | 82 | 75 | 90 | 56 | 69 | 72 | 65 | 43 | 62 | 25 | 41 |
| | MK2206 | 2 | 2 | 48 | 37 | 87 | 63 | 2 | 4 | 55 | 51 | 0 | 2 | 42 |

**Viability**

0%

10%

≥ 50%

**Figure S3**

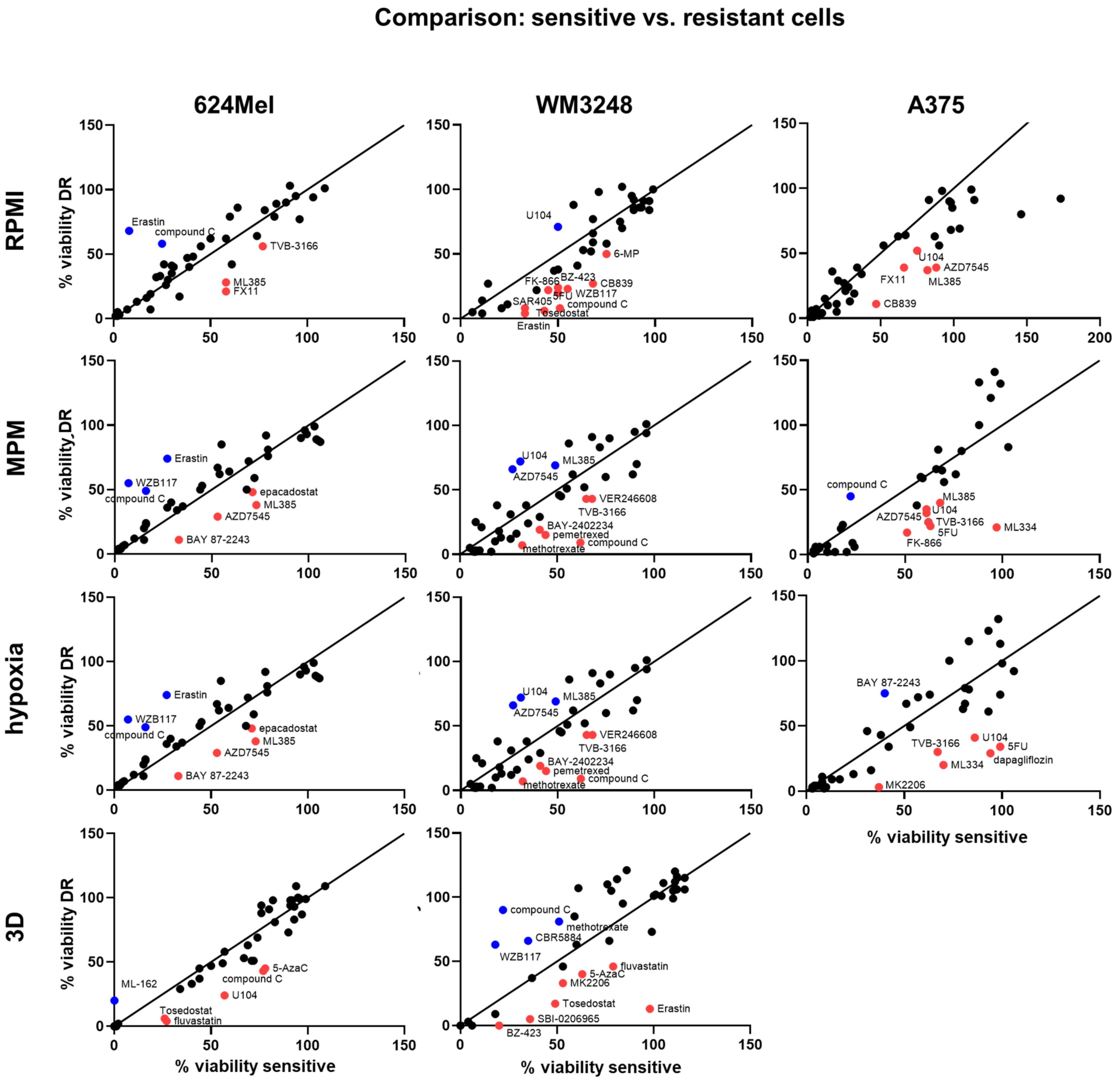

Comparison: sensitive vs. resistant cells
624Mel
WM3248
A375
RPMI
MPM
hypoxia
3D
% viability DR
% viability sensitive
Erastin
compound C
TVB-3166
ML385
FX11
U104
6-MP
FK-866
BZ-423
CB839
SAR405
5FU
WZB117
Tosedostat
AZD7545
WZB117
epacadostat
BAY 87-2243
VER246608
BAY-2402234
pemetrexed
methotrexate
ML334
FK-866
MK2206
dapagliflozin
ML-162
5-AzaC
fluvastatin
CBR5884
SBI-0206965

### Figure S4

**A** RPMI / MPM

| Drug | Row | High: 624Mel | High: 624Mel-DR | High: WM3248 | High: WM3248-DR | High: A375 | High: A375-DR | Low: 624Mel | Low: 624Mel-DR | Low: WM3248 | Low: WM3248-DR | Low: A375 | Low: A375-DR |
|---|---|---|---|---|---|---|---|---|---|---|---|---|---|
| FX11 | 1 | 35.6 | 5.6 | 16.2 | 10.8 | 24.8 | 46.7 | 1.0 | 1.0 | 1.1 | 0.9 | 1.1 | 1.3 |
| | 2 | 0.8 | 1.0 | 1.2 | 1.0 | 0.9 | 0.7 | 0.9 | 1.0 | 1.1 | 1.1 | 1.2 | 1.0 |
| | 3 | 0.9 | 1.0 | 1.0 | 0.8 | 0.9 | 0.7 | 0.9 | 0.9 | 1.1 | 1.0 | 1.1 | 1.0 |
| UK5099 | 4 | 4.2 | 3.7 | 2.2 | 2.8 | 2.6 | 15.3 | 1.1 | 1.1 | 1.0 | 1.2 | 0.9 | 1.0 |
| | 5 | 0.6 | 0.6 | 2.6 | 1.5 | 1.4 | 1.2 | 1.1 | 1.2 | 1.2 | 1.2 | 1.0 | 0.9 |
| | 6 | 1.0 | 1.1 | 0.9 | 0.5 | 0.8 | 0.5 | 1.0 | 0.9 | 1.0 | 0.1 | 1.0 | 0.2 |
| | 7 | 1.3 | 0.7 | 1.2 | 4.1 | 1.0 | 0.4 | 0.9 | 1.1 | 1.5 | 0.9 | 1.1 | 0.9 |
| | 8 | 0.7 | 0.9 | 1.5 | 2.3 | 0.7 | 0.5 | 1.0 | 1.0 | 1.1 | 0.9 | 1.0 | 1.0 |
| BZ423 | 9 | 29.1 | 18.5 | 6.4 | 9.3 | 11.5 | 16.4 | 0.9 | 1.0 | 1.2 | 1.1 | 1.1 | 1.0 |
| | 10 | 0.7 | 0.8 | 1.6 | 1.1 | 0.8 | 0.7 | 0.9 | 0.9 | 0.7 | 0.6 | 0.9 | 1.0 |
| Tosedostat | 11 | 2.1 | 2.0 | 2.6 | 3.1 | 0.9 | 2.2 | 1.2 | 1.4 | 0.9 | 1.1 | 1.7 | 1.3 |
| | 12 | 1.5 | 1.3 | 1.3 | 0.6 | 1.2 | 2.6 | 1.9 | 1.3 | 1.0 | 0.8 | 1.4 | 1.5 |
| | 13 | 0.9 | 0.9 | 1.2 | 0.8 | 2.2 | 2.1 | 0.9 | 1.1 | 0.8 | 0.9 | 1.2 | 0.9 |
| | 14 | 1.3 | 1.0 | 1.4 | 1.6 | 1.1 | 1.8 | 1.0 | 2.2 | 1.0 | 0.8 | 1.4 | 0.9 |
| | 15 | 0.9 | 1.0 | 1.1 | 0.9 | 1.2 | 0.8 | 1.0 | 1.0 | 1.1 | 1.1 | 1.0 | 1.0 |
| | 16 | 1.6 | 1.2 | 0.8 | 1.0 | 0.7 | 0.8 | 1.2 | 1.0 | 1.0 | 1.1 | 0.9 | 0.9 |
| | 17 | 0.3 | 0.9 | 0.5 | 0.1 | 0.1 | 0.0 | 0.7 | 0.9 | 0.9 | 0.3 | 0.3 | 0.8 |
| | 18 | 0.8 | 0.9 | 1.1 | 1.5 | 1.7 | 0.8 | 1.0 | 1.0 | 1.2 | 1.1 | 1.1 | 0.9 |
| | 19 | 0.6 | 0.5 | 0.8 | 2.4 | 0.8 | 0.9 | 1.2 | 1.1 | 0.1 | 0.0 | 0.6 | 0.7 |
| | 20 | 1.1 | 1.0 | 0.9 | 0.9 | 1.8 | 4.4 | 1.0 | 1.0 | 1.2 | 1.0 | 1.0 | 1.1 |
| | 21 | 0.8 | 0.7 | 1.4 | 1.0 | 1.2 | 0.9 | 1.0 | 1.5 | 2.0 | 1.4 | 0.9 | 1.1 |
| | 22 | 0.7 | 0.6 | 2.4 | 0.6 | 0.4 | 0.3 | 1.4 | 0.6 | 2.2 | 0.5 | 0.6 | 0.4 |
| | 23 | 0.6 | 0.6 | 6.5 | 3.6 | 1.8 | 0.9 | 1.1 | 0.8 | 5.7 | 2.9 | 0.9 | 1.2 |
| | 24 | 1.1 | 0.9 | 1.5 | 1.5 | 0.9 | 1.1 | 1.1 | 0.9 | 1.1 | 1.0 | 1.0 | 1.0 |
| | 25 | 3.9 | 0.5 | 1.9 | 1.4 | 1.2 | 1.3 | 1.1 | 1.1 | 1.1 | 1.0 | 1.1 | 1.0 |
| | 26 | 0.5 | 0.7 | 1.2 | 0.7 | 0.5 | 0.6 | 0.6 | 0.6 | 1.0 | 0.6 | 0.7 | 0.3 |
| | 27 | 0.3 | 0.4 | 0.8 | 0.7 | 0.3 | 0.3 | 0.3 | 0.6 | 0.7 | 0.7 | 0.3 | 0.2 |
| | 28 | 0.8 | 0.8 | 1.7 | 3.1 | 1.4 | 1.2 | 0.8 | 0.9 | 2.0 | 2.8 | 1.0 | 1.2 |
| | 29 | 0.5 | 0.5 | 1.9 | 6.2 | 2.1 | 2.7 | 0.9 | 0.9 | 1.4 | 3.6 | 1.7 | 3.7 |
| | 30 | 0.9 | 0.9 | 1.5 | 3.4 | 2.6 | 4.2 | 1.0 | 0.9 | 1.1 | 2.5 | 3.6 | 7.6 |
| | 31 | 1.8 | 1.3 | 2.0 | 1.2 | 1.0 | 1.5 | 1.2 | 1.5 | 1.0 | 1.2 | 1.1 | 1.6 |
| | 32 | 1.5 | 1.4 | 1.1 | 0.8 | 0.8 | 1.8 | 0.9 | 1.1 | 0.9 | 0.6 | 0.9 | 1.1 |
| | 33 | 0.8 | 1.3 | 1.2 | 1.2 | 0.9 | 1.0 | 1.2 | 1.0 | 1.1 | 1.0 | 1.0 | 0.9 |
| | 34 | 0.5 | 0.8 | 1.5 | 2.1 | 1.1 | 1.1 | 1.0 | 1.1 | 0.6 | 0.3 | 0.8 | 1.0 |
| | 35 | 0.6 | 0.8 | 1.6 | 1.0 | 1.2 | 1.5 | 1.0 | 1.1 | 1.1 | 1.0 | 1.1 | 0.8 |
| | 36 | 0.8 | 1.0 | 1.6 | 1.0 | 1.3 | 1.1 | 1.0 | 1.1 | 1.2 | 0.9 | 1.1 | 1.0 |
| | 37 | 1.1 | 1.1 | 1.1 | 0.8 | 1.4 | 1.3 | 1.0 | 1.1 | 1.2 | 0.9 | 1.1 | 0.9 |
| | 38 | 1.1 | 1.1 | 1.2 | 1.1 | 1.7 | 1.5 | 1.0 | 1.1 | 1.2 | 1.1 | 1.1 | 0.9 |
| | 39 | 1.0 | 1.0 | 1.8 | 2.0 | 1.5 | 1.2 | 1.4 | 1.3 | 1.4 | 1.5 | 1.6 | 1.6 |
| SB204990 | 40 | 1.8 | 1.2 | 2.8 | 2.3 | 2.0 | 2.4 | 1.0 | 1.1 | 1.2 | 1.1 | 0.9 | 0.9 |
| | 41 | 1.1 | 0.8 | 1.3 | 1.7 | 1.5 | 2.2 | 1.3 | 1.3 | 1.2 | 1.9 | 1.4 | 2.0 |
| | 42 | 0.6 | 0.6 | 0.9 | 0.7 | 4.4 | 38.9 | 1.3 | 1.2 | 0.8 | 0.6 | 0.8 | 1.0 |

**B** MPM / RPMI

| Drug | Row | High: 624Mel | High: 624Mel-DR | High: WM3248 | High: WM3248-DR | High: A375 | High: A375-DR | Low: 624Mel | Low: 624Mel-DR | Low: WM3248 | Low: WM3248-DR | Low: A375 | Low: A375-DR |
|---|---|---|---|---|---|---|---|---|---|---|---|---|---|
| | 1 | 0.0 | 0.2 | 0.1 | 0.1 | 0.0 | 0.0 | 1.0 | 1.0 | 0.9 | 1.1 | 0.9 | 0.8 |
| | 2 | 1.3 | 1.0 | 0.8 | 1.0 | 1.1 | 1.5 | 1.1 | 1.0 | 1.0 | 0.9 | 0.8 | 1.0 |
| | 3 | 1.1 | 1.0 | 1.0 | 1.2 | 1.1 | 1.3 | 1.1 | 1.1 | 0.9 | 1.0 | 0.9 | 1.0 |
| | 4 | 0.2 | 0.3 | 0.5 | 0.4 | 0.4 | 0.1 | 0.9 | 0.9 | 1.0 | 0.8 | 1.1 | 1.0 |
| | 5 | 1.6 | 1.7 | 0.4 | 0.7 | 0.7 | 0.8 | 0.9 | 0.9 | 0.8 | 0.8 | 1.0 | 1.1 |
| CB839 | 6 | 1.0 | 0.9 | 1.1 | 2.2 | 1.3 | 2.2 | 1.0 | 1.1 | 1.0 | 9.3 | 1.0 | 6.1 |
| | 7 | 0.8 | 1.4 | 0.8 | 0.2 | 1.0 | 2.2 | 1.1 | 0.9 | 0.7 | 1.2 | 0.9 | 1.1 |
| | 8 | 1.3 | 1.1 | 0.7 | 0.4 | 1.4 | 2.2 | 1.0 | 1.0 | 0.9 | 1.1 | 1.0 | 1.0 |
| | 9 | 0.0 | 0.1 | 0.2 | 0.1 | 0.1 | 0.1 | 1.1 | 1.0 | 0.8 | 0.9 | 0.9 | 1.0 |
| | 10 | 1.5 | 1.3 | 0.6 | 0.9 | 1.3 | 1.4 | 1.2 | 1.1 | 1.5 | 1.6 | 1.2 | 1.0 |
| | 11 | 0.5 | 0.5 | 0.4 | 0.3 | 1.1 | 0.5 | 0.8 | 0.7 | 1.2 | 0.9 | 0.6 | 0.8 |
| | 12 | 0.7 | 0.7 | 0.8 | 1.6 | 0.8 | 0.4 | 0.5 | 0.8 | 1.0 | 1.3 | 0.7 | 0.7 |
| | 13 | 1.1 | 1.1 | 0.8 | 1.2 | 0.5 | 0.5 | 1.1 | 0.9 | 1.2 | 1.1 | 0.9 | 1.1 |
| | 14 | 0.7 | 1.0 | 0.7 | 0.6 | 0.9 | 0.6 | 1.0 | 0.4 | 1.0 | 1.2 | 0.7 | 1.2 |
| | 15 | 1.1 | 1.0 | 0.9 | 1.1 | 0.8 | 1.2 | 1.0 | 1.0 | 0.9 | 0.9 | 1.0 | 1.0 |
| | 16 | 0.6 | 0.9 | 1.2 | 1.0 | 1.3 | 1.3 | 0.8 | 1.0 | 1.0 | 0.9 | 1.1 | 1.1 |
| Erastin | 17 | 3.2 | 1.1 | 1.9 | 13.0 | 6.9 | 32.3 | 1.3 | 1.1 | 1.1 | 3.8 | 3.9 | 1.3 |
| | 18 | 1.2 | 1.1 | 0.9 | 0.7 | 0.6 | 1.3 | 1.0 | 1.0 | 0.8 | 0.9 | 0.9 | 1.1 |
| ML162 | 19 | 1.7 | 2.0 | 1.2 | 0.4 | 1.3 | 1.1 | 0.8 | 0.9 | 11.9 | 29.6 | 1.7 | 1.4 |
| | 20 | 0.9 | 1.0 | 1.1 | 1.1 | 0.6 | 0.2 | 1.0 | 1.0 | 0.9 | 1.0 | 1.0 | 0.9 |
| | 21 | 1.3 | 1.4 | 0.7 | 1.1 | 0.8 | 1.1 | 1.0 | 0.7 | 0.5 | 0.7 | 1.1 | 0.9 |
| FK866 | 22 | 1.4 | 1.8 | 0.4 | 1.7 | 2.5 | 3.4 | 0.7 | 1.6 | 0.5 | 1.9 | 1.6 | 2.6 |
| | 23 | 1.7 | 1.7 | 0.2 | 0.3 | 0.5 | 1.1 | 0.9 | 1.3 | 0.2 | 0.3 | 1.1 | 0.8 |
| | 24 | 0.9 | 1.1 | 0.7 | 0.7 | 1.1 | 0.9 | 1.0 | 1.1 | 0.9 | 1.0 | 1.0 | 1.0 |
| | 25 | 0.3 | 2.1 | 0.5 | 0.7 | 0.9 | 0.8 | 1.0 | 0.9 | 0.9 | 1.0 | 0.9 | 1.0 |
| 5FU | 26 | 2.1 | 1.5 | 0.8 | 1.4 | 2.2 | 1.7 | 1.7 | 1.6 | 1.0 | 1.7 | 1.4 | 3.0 |
| 6MP | 27 | 3.6 | 2.5 | 1.2 | 1.4 | 3.0 | 3.7 | 2.9 | 1.6 | 1.4 | 1.5 | 3.3 | 4.6 |
| | 28 | 1.3 | 1.3 | 0.6 | 0.3 | 0.7 | 0.8 | 1.2 | 1.1 | 0.5 | 0.4 | 1.0 | 0.9 |
| | 29 | 1.9 | 2.0 | 0.5 | 0.2 | 0.5 | 0.4 | 1.2 | 1.2 | 0.7 | 0.3 | 0.6 | 0.3 |
| | 30 | 1.2 | 1.1 | 0.7 | 0.3 | 0.4 | 0.2 | 1.0 | 1.1 | 0.9 | 0.4 | 0.3 | 0.1 |
| | 31 | 0.6 | 0.8 | 0.5 | 0.8 | 1.0 | 0.7 | 0.8 | 0.7 | 1.0 | 0.8 | 0.9 | 0.6 |
| | 32 | 0.7 | 0.7 | 0.9 | 1.2 | 1.2 | 0.6 | 1.1 | 0.9 | 1.2 | 1.6 | 1.1 | 0.9 |
| | 33 | 1.2 | 0.8 | 0.8 | 0.8 | 1.1 | 1.0 | 0.9 | 1.0 | 0.9 | 1.0 | 1.0 | 1.1 |
| | 34 | 1.9 | 1.2 | 0.7 | 0.5 | 1.0 | 0.9 | 1.1 | 0.9 | 1.5 | 2.9 | 1.2 | 1.0 |
| | 35 | 1.7 | 1.3 | 0.6 | 1.0 | 0.8 | 0.7 | 1.0 | 0.9 | 0.9 | 1.0 | 0.9 | 1.3 |
| | 36 | 1.2 | 1.0 | 0.6 | 1.0 | 0.8 | 0.9 | 1.0 | 0.9 | 0.8 | 1.1 | 0.9 | 1.0 |
| | 37 | 0.9 | 0.9 | 0.9 | 1.2 | 0.7 | 0.8 | 1.0 | 0.9 | 0.9 | 1.1 | 0.9 | 1.1 |
| | 38 | 0.9 | 0.9 | 0.8 | 0.9 | 0.6 | 0.7 | 1.0 | 0.9 | 0.8 | 0.9 | 0.9 | 1.1 |
| | 39 | 1.0 | 1.0 | 0.6 | 0.5 | 0.7 | 0.8 | 0.7 | 0.8 | 0.7 | 0.7 | 0.6 | 0.6 |
| | 40 | 0.6 | 0.9 | 0.4 | 0.4 | 0.5 | 0.4 | 1.0 | 0.9 | 0.9 | 0.9 | 1.1 | 1.2 |
| | 41 | 0.9 | 1.3 | 0.8 | 0.6 | 0.7 | 0.4 | 0.8 | 0.8 | 0.8 | 0.5 | 0.7 | 0.5 |
| | 42 | 1.6 | 1.7 | 1.2 | 1.4 | 0.0 | 0.0 | 0.8 | 0.8 | 1.2 | 1.5 | 1.2 | 1.0 |

Ratio: >10 / 2.5 / 1

#### Figure S5

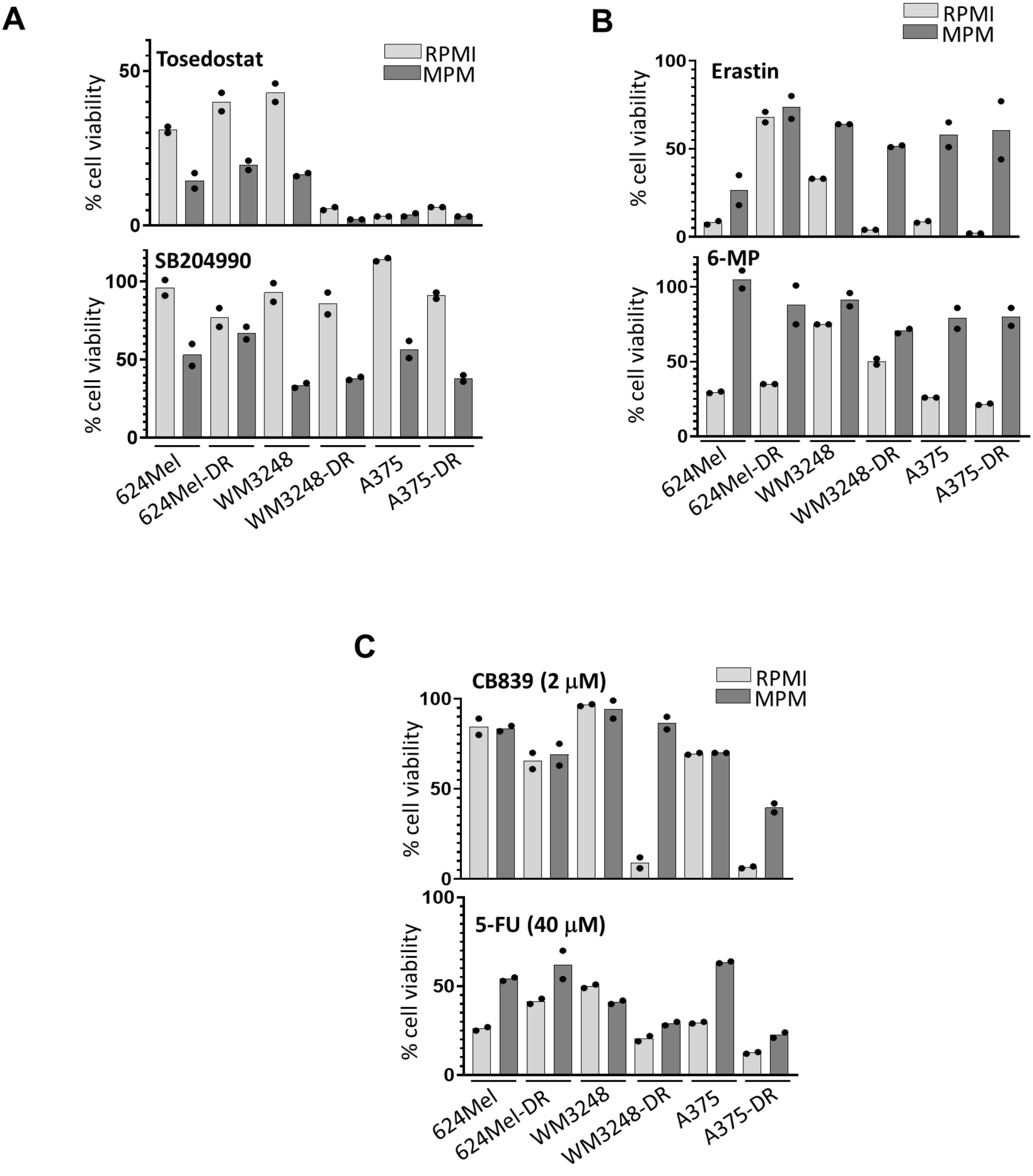

A
Tosedostat
SB204990
RPMI
MPM
% cell viability
B
Erastin
6-MP
C
CB839 (2 μM)
5-FU (40 μM)
624Mel
624Mel-DR
WM3248
WM3248-DR
A375
A375-DR

#### Figure S6

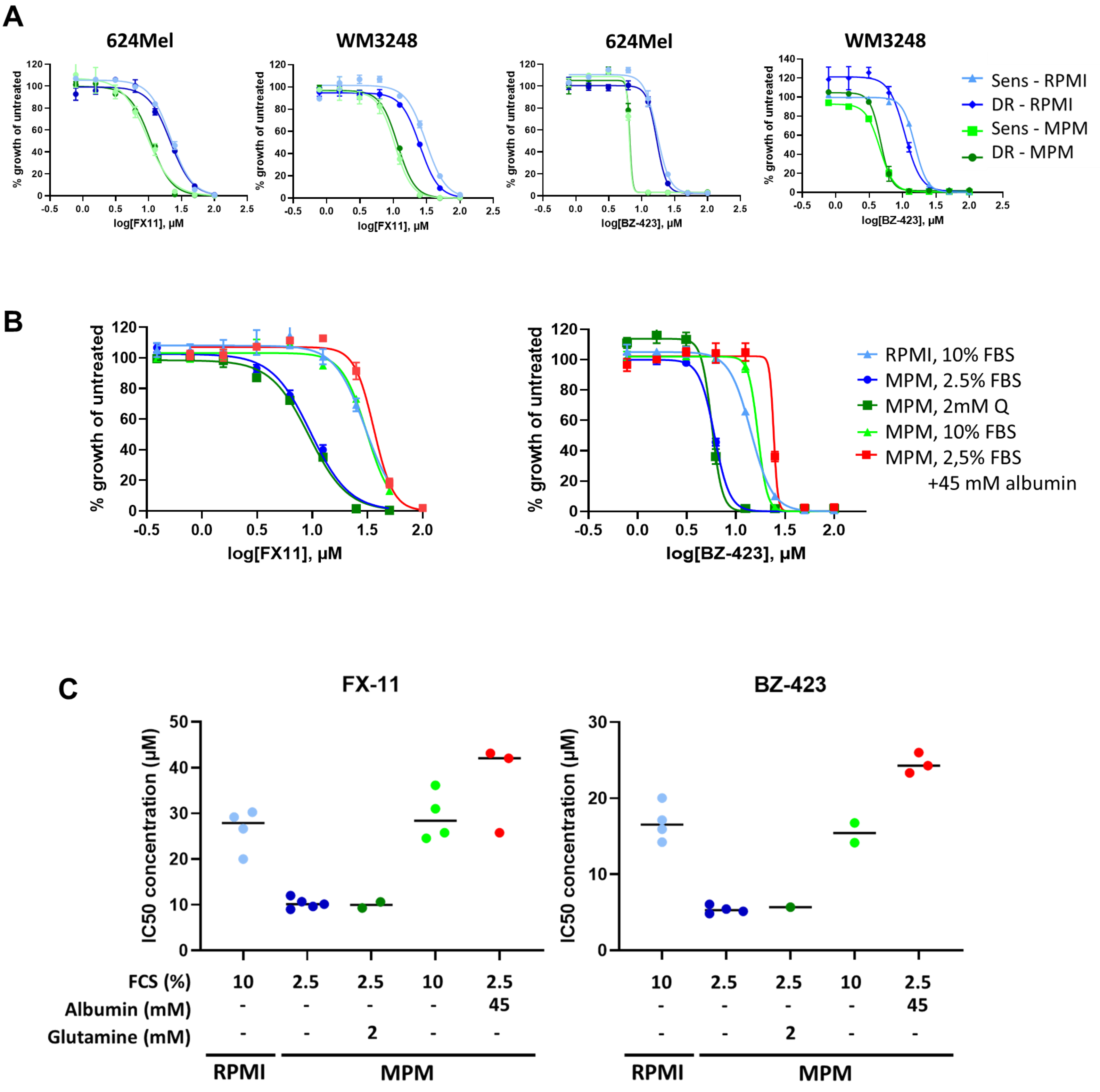
A
624Mel
WM3248
624Mel
WM3248
% growth of untreated
log[FX11], μM
log[BZ-423], μM
Sens - RPMI
DR - RPMI
Sens - MPM
DR - MPM
B
% growth of untreated
log[FX11], μM
log[BZ-423], μM
RPMI, 10% FBS
MPM, 2.5% FBS
MPM, 2mM Q
MPM, 10% FBS
MPM, 2,5% FBS
+45 mM albumin
C
FX-11
BZ-423
IC50 concentration (μM)
FCS (%)
Albumin (mM)
Glutamine (mM)
10
2.5
2.5
10
2.5
45
2
RPMI
MPM

#### Figure S7

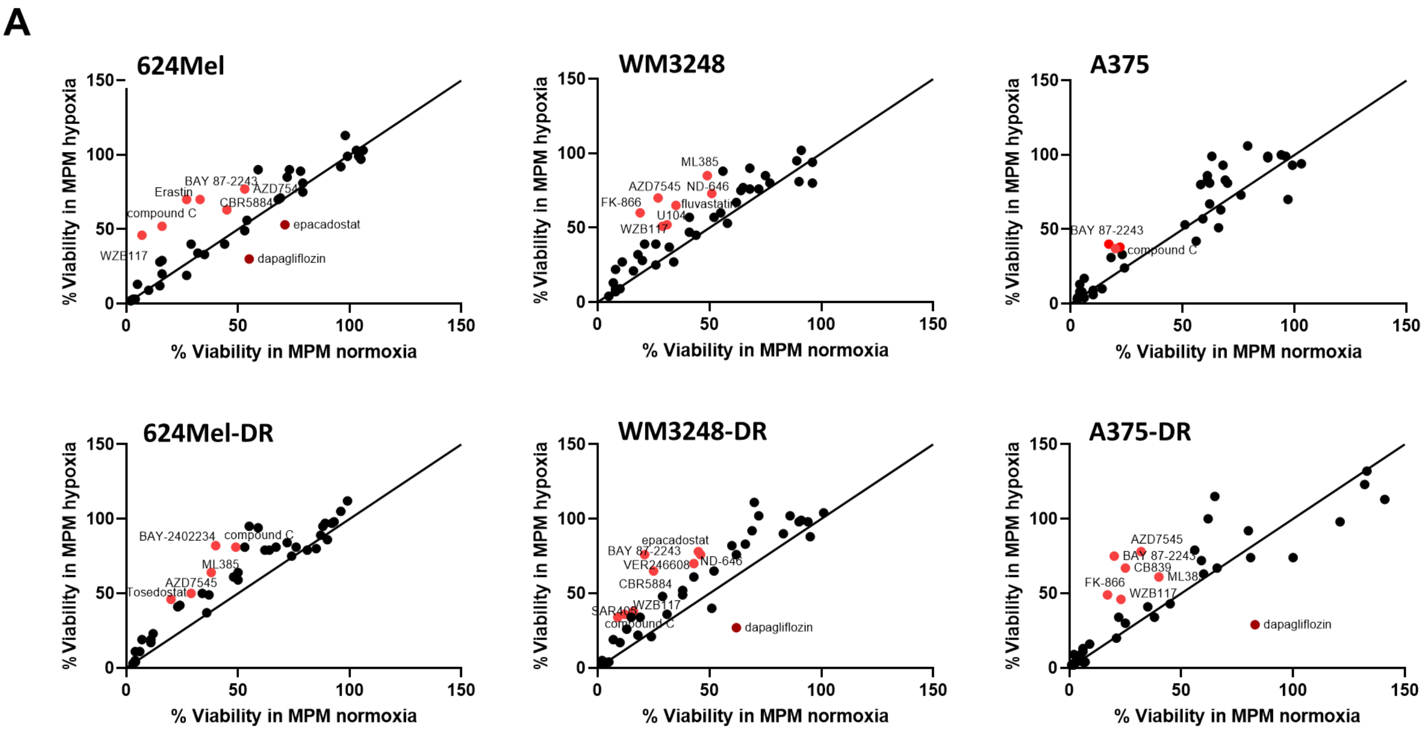
A
624Mel
WM3248
A375
624Mel-DR
WM3248-DR
A375-DR
% Viability in MPM hypoxia
% Viability in MPM normoxia
BAY 87-2243
Erastin
AZD7545
CBR5884
compound C
epacadostat
WZB117
dapagliflozin
ML385
ND-646
FK-866
fluvastatin
U104
BAY-2402234
Tosedostat
VER246608
SAR405
CB839

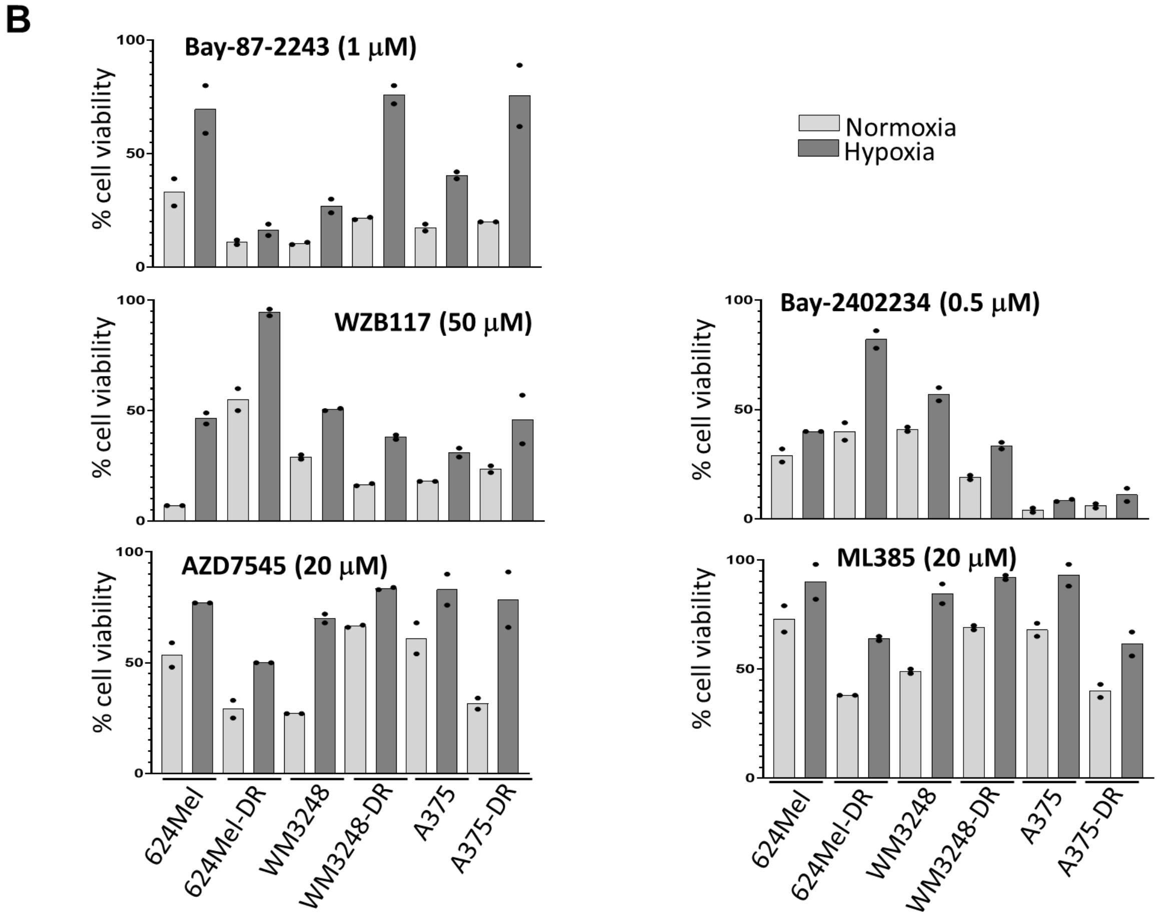
B
Bay-87-2243 (1 μM)
WZB117 (50 μM)
Bay-2402234 (0.5 μM)
AZD7545 (20 μM)
ML385 (20 μM)
% cell viability
Normoxia
Hypoxia
624Mel
624Mel-DR
WM3248
WM3248-DR
A375
A375-DR
